# Detection of Maximal Exact Matches in Terabases of Genome Assemblies in Compressed Space and Sublinear Time

**DOI:** 10.64898/2026.11.10.765459

**Authors:** Ahsan Sanaullah, Degui Zhi, Heng Li, Shaojie Zhang

## Abstract

Detection of long exact matches between a pair of genomes has historically been a fundamental operation in genomics. Similarly, detection of long exact matches between all pairs of haplotypes in a large panel of genotyping array data or whole-genome sequencing variant calls is a fundamental operation in population genetics. Thanks to various recent pangenome efforts, for the first time, large panels of high-quality genome assemblies are being constructed. Therefore, we propose the intersection of these two approaches: detection of long exact matches between all pairs of assemblies in panels of hundreds to thousands of high-quality assemblies. Since these datasets would contain trillions of basepairs, practically detecting all exact matches requires algorithms operating in at most linear time and compressed space. Previous methods such as Mumemto achieve this, but running them on terabase-sized pangenomes still took very large space and long time. Here, we describe more efficient algorithms for outputting all long exact matches between all pairs of assemblies in compressed space and sublinear time given a compressed index of the panel. Given the construction of an *O*(*r*) space compressed index—a one-time cost that can be done in compressed space and close to linear time with previous work—we describe three algorithms taking *O*(*r*) words of memory: (1) computing all supermaximal exact matches (SMEMs) in *O*(*r*) time, (2) computing all supermaximal exact matches and their occurrences in *O*(*r* + *occ*) time, and (3) computing all maximal exact matches (MEMs) in *O*(*r* + *occ*) time, where *r* is the number of runs in the Burrows-Wheeler transform of the panel of assemblies and *occ* is the number of occurrences or matches outputted. Our algorithms use 7.5 times less memory than state-of-the-art competitors on a dataset of human centromere assemblies from 625 haplotypes. They are also 20 times faster if index construction time is counted, and 2,500 times faster if index construction is excluded. Finally, we perform an analysis of all-vs-all MEMs in all well-assembled centromeres of the Human Pangenome Reference Consortium Release 2 dataset that reveals multiple novel insights into structural variation in centromeres. For example, we find that the active *α*-satellite higher-order repeat (HOR) array of Chromosome 2 is flanked by a *∼* 200 kbp inversion on one side and a *∼* 200–300 kbp pseudopalindromic sequence on the other. We also detect large segmental duplications in the centromere of every chromosome except Chromosome Y. These segmental duplications range in size from hundreds of kbp to multiple Mbp and occur both in and out of the active *α*-satellite HOR arrays. Our computational results make pairwise MEM analyses possible in pangenomes, while our biological results show they are likely to be useful. Our algorithms are implemented as TeraMEM in the TeraTools suite available at https://github.com/ucfcbb/TeraTools.

## 1 Introduction

A central problem in bioinformatics is to find matches in biological sequences. In traditional genomics, these long matches are called Maximal Exact Matches (MEM) or sometimes anchors in the alignment literature and are utilized by widely used genome-to-genome alignment [21] and read-to-genome alignment [22] software. In population genetics, these long matches are typically just called long matches and come with the additional restriction that the matches must occur in the same position in multiple strings [12].

Historically, there have been a low number of high-quality human genome assemblies and a high number of haplotypes with variant calls. Therefore, the algorithms computing long matches in these two fields diverged. Population geneticists developed efficient algorithms for outputting all pairwise matches in thousands to millions of haplotypes [12], while genomicists lack this capability for even hundreds of assemblies. Hundreds of high-quality human genomes have been assembled in the past few years due to increasing quality and decreasing cost of long read sequencing. Multiple groups [23, 24, 26] are generating these large human pangenomes, and current plans aim to increase their size to thousands of high-quality assemblies in the next few years (we define a pangenome as a set of high-quality assemblies from now on). Therefore, we aim to provide the pairwise matching capability available in variant data in large pangenomes as well. A major hurdle in doing so is that for the same number of haplotypes, a pangenome contains orders of magnitude more raw data than a panel of haplotype variant calls, around three hundred to three thousand times more.

We address this challenge by developing algorithms running in time and space sublinear to the size of the dataset given a compressed index of the pangenome. Even for only 230 genomes (the current Human Pangenome Reference Consortium (HPRC) release [24]), there is no method capable of efficiently finding all pairwise long enough exact matches between assemblies. Mumemto [37] comes the closest; it uses compressed space and a parallelized version of it [36] can output multi-maximal unique matches (multi-MUMs), a special case of match that occurs in every genome exactly once. It can also output all-vs-all pairwise matches (partial multi-MEMs), but this mode is sequential only. Its runtime is linear to the size of the pangenome and its memory usage is proportional to the size of a grammar of the pangenome, *PFP* [5]. For the HPRC, Mumemto requires multiple terabytes of RAM and possibly a week of run time to compute all pairwise MEMs. In this work, we develop algorithms for outputting all pairwise MEMs in both sublinear time and compressed space. In particular, our algorithms run in time and space linear in *r* and *occ*, where *r* is the number of runs in the Burrows-Wheeler transform and *occ* is the number of matches outputted. *r* has been shown to be much smaller than |*PFP*| in practice and up to 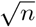 times smaller than it in theory, where *n* is the total number of basepairs in the pangenome [18, 27]. Our implementation, TeraMEM, is up to 20 times faster and uses 7.5 times less space than Mumemto if compressed index construction costs are included for both tools and is more than 2,500 times faster if index construction is excluded for both tools.

Finally, we perform a MEM-based preliminary biological analysis to demonstrate the utility of pairwise MEM-based analyses in pangenomes. We use our algorithms to call all-vs-all MEMs in a dataset of well-assembled centromeres from 453 human haplotypes, annotated by Gao et al. [16]. Next, we filter these MEMs to cases that may indicate structural variation. In doing so, we recover known relationships: an inversion larger than a million basepairs in the Chromosome 1 centromere [1] and interchromosomal exchange of whole centromeres between Chromosomes 13 and 21 [14]. In addition, we possibly discover previously unknown variation in the centromeres: inverted repeats and pseudopalindromes up to hundreds of kilobases long just outside of the active *α*-satellite higher-order repeat (HOR) arrays in Chromosomes 2, 4, 7, 11, 20, 22, and X, and segmental duplications hundreds of kilobases to megabases long in centromeres of all chromosomes except Chromosome Y. Further analysis is needed to confirm the novel results. If confirmed, they may imply that large segmental duplication in centromeres is frequent enough to play an active role in their evolution. In summary, our contributions are as follows (all algorithms require an *O*(*r*) space index as input and *occ* is the number of occurrences or MEMs outputted):

- Theorem 1: An *O*(*r*) time and space algorithm for computing all supermaximal exact matches (SMEM).
- Theorem 2: An *O*(*r* +*occ*) time and *O*(*r*) space algorithm computing all SMEMs and their occurrences.
- Theorem 3: An *O*(*r* + *occ*) time and *O*(*r*) space algorithm for computing all MEMs.
- Section 4.2: Discovering possibly new peri/centromeric variation with a pairwise MEM-based analysis: segmental duplications in and inversions around the active *α*-satellite HOR arrays.

## 2 Background

We define a text *T* as a string of length *n*, composed of characters from an alphabet Σ = {0, 1, 2, …, *σ –* 1}. The character 0 occurs exactly once in *T*, at the end. We use |*-*| to denote the length of a string or the size of a set; then, let |Σ| = *σ* and |*T*| = *n*. Strings and arrays are 0-indexed and the concatenation of *p* and *q* is denoted by *pq*; then *T* = *T* [0]*T* [1]*T* [2] … *T* [*n −* 1]. For 0 *≤ i ≤ j < n, T* [*i, j*] denotes *T* [*i*]*T* [*i* + 1]*T* [*i* + 2] … *T* [*j*]. *T* [*a, a −* 1] denotes the empty string, *ε*. Suffix *i* of *T* is *T* [*i, n −* 1]. Let *occ*(*P, T*) be the number of times *P* occurs as a substring in *T* . The length of the longest common prefix of strings *p* and *q* is *lcp*_*l*_(*p, q*) = *max{i ∈* [0, min(|*p*|, |*q*|)]: *p*[0, *i −* 1] = *q*[0, *i −* 1]*}*. The longest common prefix of *p* and *q* is *lcp*(*p, q*) = *p*[0, *lcp*_*l*_(*p, q*) *−* 1] = *q*[0, *lcp*_*l*_(*p, q*) *−* 1]. For convenience, we define some slight abuses of notation: *T* [*−*1] = *T* [*n*] = *ε, T* [*i, j*] = *ε* if *i* = *−*1 or *j* = *n*, and *occ*(*ε, T*) = 1.

### 2.1 Suffix Array and Burrows-Wheeler transform

The suffix array, *SA*, of a text *T* is a permutation of [0, *n −* 1] such that *T* [*SA*[*i*], *n −* 1] is the *i*-th lexi-cographically smallest suffix of *T* . The longest common prefix array of *T, LCP*, is an array of length *n* that stores the length of the longest common prefix between adjacent suffixes in the suffix array, *LCP* [*i*] = *lcp*_*l*_(*T* [*SA*[*i*], *n −* 1], *T* [*SA*[*I −* 1], *n −* 1]). Note, *LCP* [0] = 0. Let *LCP*_*d*_[*i*] = *LCP* [(*i* + 1) mod *n*].

The *ϕ* array is a permutation of [0, *n −* 1] storing at position *i* the starting index of the lexicographically largest suffix of *T* lexicographically smaller than *T* [*i, n −* 1]. In other words, if *SA*[*j*] = *i*, then *ϕ*[*i*] = *SA*[*j −* 1]. Note if *j* = 0, we define *ϕ*[*i*] = *ϕ*[*SA*[0]] = *SA*[*n −* 1]. The *ϕ*^*−*1^ array is the inverse of *ϕ*: for *SA*[*j*] = *i, ϕ*^*−*1^[*i*] = *SA*[*j* + 1] and *ϕ*^*−*1^[*SA*[*n −* 1]] = *SA*[0].

The *P LCP*_*u*_ array stores at position *i* the length of the longest common prefix between *T* [*i, n −* 1] and *T* [*ϕ*[*i*], *n −* 1]. Similarly, the *P LCP*_*d*_ array stores at position *i* the length of the longest common prefix between *T* [*i, n −* 1] and *T* [*ϕ*^*−*1^[*i*], *n −* 1]. Then the following equivalencies hold: *P LCP*_*u*_[*SA*[*i*]] = *LCP* [*i*] and *P LCP*_*d*_[*SA*[*i*]] = *LCP* [(*i* + 1) mod *n*].

The Burrows-Wheeler transform of *T, BWT*, is a string of length *n* storing in position *i* the character preceding the *i*-th lexicographically smallest suffix. Then, *BWT* [*i*] = *T* [(*SA*[*i*] *−* 1) mod *n*]. Therefore, the *BWT* of *T* is a permutation of *T* .

### 2.2 Run-Length Encoding and Move Structure

A run in the *BWT* is a range, [*i, j*], such that *BWT* [*i −* 1] ≠ *BWT* [*i*], *BWT* [*j* + 1] ≠ *BWT* [*j*], and *∀k ∈* [*i, j*], *BWT* [*k*] = *BWT* [*i*]. Let *r* be the number of runs in the *BWT* . The *k*-th *BWT* run is denoted by *r*_*k*_, and if *r*_*k*_ = [*i, j*], then its start is *r*_*k*_.*s* = *i* and its end is *r*_*k*_.*e* = *j*. Then, *r*_0_.*s* = 0 and *r*_*r −* 1_.*e* = *n −* 1. Let *r*_*k*_.*c* store the character of the *k*-th run, *r*_*k*_.*c* = *BWT* [*r*_*k*_.*s*]. Then the run-length encoded *BWT* (RLBWT) can be fully specified by *n* and all *r*_*k*_.*s* and *r*_*k*_.*c* values.

The move structure is a data structure introduced by Nishimoto and Tabei that compresses “runny” permutations while retaining constant time evaluation [28]. Let *f* be a permutation of [0, *n −* 1]; we call [*i, j*] a “conserved interval” in *f* if *∀k ∈* [*i, j*], *f* (*k*) = *f* (*i*) + (*k − i*). We may represent the range [0, *n −* 1] as a sequence of disjoint conserved intervals fully covering [0, *n −* 1]. We will represent a sequence of disjoint intervals covering [0, *n −* 1] as the sorted sequence of their starting positions along with *n*. Let *I* be this representation, *I* = {*i*_0_ = 0, *i*_1_, *i*_2_, …, *i* _|*I*|*−* 2_, *i* _|*I*|*−* 1_ = *n*}. Then there are |*I*| *−* 1 conserved intervals and the *j*-th conserved interval is [*i*_*j*_, *i*_*j*+1_ 1] for *j ∈* [0, |*I* |*−* 2]. For all *k ∈* [0, *n −* 1], given *x* where *i*_*x*_ *≤ k < i*_*x*+1_, the move structure allows computing *f* (*k*) and *x*^*′*^ where 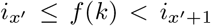 in constant time and *O*(|*I* |) space^1^. See [28, 6] for the proofs for the time and space complexities of the move structure and [28, 3, 7] for its construction algorithms.

#### Claim 1

*Let* Φ(*i*) = |{*j ≤ i*: *SA*[*r*_*k*_.*s*] = *j* for some *k ∈* [0, *r −* 1]}|*−* 1. *There exists an O*(*r*) *space data structure that computes ϕ*[*i*], Φ(*ϕ*[*i*]), *and P LCP*_*u*_[*i*] *in constant time given* Φ(*i*).

*Proof*. See proof in Appendix A.

#### Claim 2

*Let* Φ_*inv*_(*i*) = |{*j ≤ i*: *SA*[*r*_*k*_.*e*] = *j* for some *k ∈* [0, *r −* 1]}| *−* 1. *There exists an O*(*r*) *space data structure that computes ϕ*^*−*1^[*i*], Φ_*inv*_(*ϕ*^*−*1^[*i*]), *and P LCP*_*d*_[*i*] *in constant time given* Φ_*inv*_(*i*).

*Proof*. See proof in Appendix A.

We will use square brackets for *ϕ, ϕ*^*−*1^, *P LCP*_*u*_,and *P LCP*_*d*_ when referring to a value directly. If we instead use parentheses, this implies that the value is computed by the structure of Claim 1 in constant time and *O*(*r*) space for *ϕ* and *P LCP*_*u*_ (or the structure of Claim 2 for *ϕ*^*−*1^ and *P LCP*_*d*_). Therefore *ϕ*[*i*] refers to the value contained at the *i*-th position in a hypothetical *ϕ* array of length *n*, while *ϕ*(*i*) implies Φ(*i*) is known and (*ϕ*[*i*], Φ(*ϕ*[*i*])) is computed in constant time and *O*(*r*) space using the *ϕ* move structure and Φ(*i*).

## 3 Methods

A maximal exact match (MEM) in *T* is defined by two intervals on *T*: call them [*i, j*] and [*i*^*′*^, *j*^*′*^]. *T* [*i, j*] = *T* [*i*^*′*^, *j*^*′*^] is a MEM iff *i ≠ i*^*′*^, *T* [*i −* 1] *T* [*i*^*′*^ *−* 1], and *T* [*j* + 1] *T* [*j*^*′*^ + 1]. An interval [*i, j*] of *T* is a supermaximal exact match (SMEM) iff *occ*(*T* [*i, j*], *T*) *>* 1, *occ*(*T* [*i −* 1, *j*], *T*) = 1, and *occ*(*T* [*i, j* + 1], *T*) = 1.

We define *LCP*_*r*_, an *O*(*r*) space data structure built on *T* . *LCP*_*r*_ allows the computation of *ϕ, ϕ*^*−*1^, *P LCP*_*u*_, and *P LCP*_*d*_ in constant time. It has the following components.

- *I*^*u*^, *RP LCP*_*u*_, and the *ϕ* move data structure as described in the proof of Claim 1.
- *I*^*d*^, *RP LCP*_*d*_, and the *ϕ*^*−*1^ move data structure as described in the proof of Claim 2.
- An array *RL*^*u*^ of length *r* that maps suffixes at the start of maximal runs in the *BWT* to the *ϕ* conserved intervals that they start 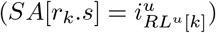.
- An array *RL*^*d*^ of length *r* that maps suffixes at the end of maximal runs in the *BWT* to the *ϕ*^*−*1^ conserved intervals that they start 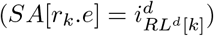.

In Section 3.1, we describe two algorithms given an *LCP*_*r*_ of the text: one outputting all SMEMs in *O*(*r*) time and space, and one outputting all SMEMs and their occurrences in *O*(*r*) space and *O*(*r* + *occ*) time. In Section 3.2, we describe an algorithm for outputting all MEMs length *L* or longer of a text in *O*(*r* + *occ*) time and *O*(*r*) space given an *LCP*_*r*_ of the text, where *L* is independent of the index and set at algorithm run time. *occ* is the number of MEMs outputted or SMEM occurrences found.

### 3.1 Supermaximal Exact Matches Algorithm

The algorithm we present here relies on two simple properties of SMEMs:

#### Lemma 1

*For any SMEM T* [*i, j*], *i occurs in the suffix array at the top or bottom of a maximal run in the BWT (∃k ∈* [0, *r −* 1] s.t. *SA*[*r*_*k*_.*s*] = *i* or *SA*[*r*_*k*_.*e*] = *i)*.

*Proof*. The definition of SMEM implies *occ*(*T* [*i, j*], *T*) *>* 1. Therefore, there is at least one suffix of *T* other than *T* [*i, n −* 1] that has *T* [*i, j*] as a prefix, call it *T* [*i*^*′*^, *n −* 1]. By the lexicographic ordering of suffixes in the suffix array and the fact that *occ*(*T* [*i, j* + 1], *T*) = 1, one such *i*^*′*^ is adjacent to *i* in the suffix array. Let *SA*[*ι*] = *i*. If *SA*[*ι* + 1] = *i*^*′*^, then *BWT* [*ι*]*≠ BWT* [*ι* + 1] by *i* = 0 *∨ occ*(*T* [*i −* 1, *j*], *T*) = 1 and *ι* is the bottom of a maximal run in the *BWT* . Otherwise, if *SA*[*ι −* 1] = *i*^*′*^, then *BWT* [*ι*]*≠ BWT* [*ι −* 1] by *i* = 0 *∨ occ*(*T* [*i −* 1, *j*], *T*) = 1 and *ι* is the top of a maximal run in the *BWT* .

#### Lemma 2

*T* [*i, j*] *is an SMEM if and only if* max(*PLCP*_*u*_[*i−*1], *P LCP*_*d*_[*i−*1]) *≤* max(*P LCP*_*u*_[*i*], *P LCP*_*d*_[*i*]) *and j* = *i* + max(*P LCP*_*u*_[*i*], *P LCP*_*d*_[*i*]) *−* 1.

*Proof*. See proof in Appendix A.

Given *LCP*_*r*_ of a text *T*, Lemmas 1 and 2 enable an algorithm for computing all SMEMs of *T* in *O*(*r*) space and time. This is achieved by checking if max(*P LCP*_*u*_(*i −* 1), *P LCP*_*d*_(*i −* 1)) *≤* max(*P LCP*_*u*_(*i*), *PLCP*_*d*_(*i*)) for each suffix *i* that occurs at the top or bottom of a maximal run in the *BWT*, if so, report *T* [*i, i* + max(*P LCP*_*u*_(*i*), *P LCP*_*d*_(*i*)) *−* 1] as an SMEM. All suffixes *i* occurring at the top or bottom of a maximal run in the *BWT* exist as either 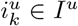 or 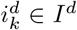 respectively for some *k ∈* [0, *r −* 1]. All *P LCP*_*u*_ and *P LCP*_*d*_ values are recovered in constant time through the structures described in Claim 1 and Claim 2 respectively. This algorithm is depicted in Figure 1. High-level pseudocode of this algorithm can be seen in Algorithm 1 in Appendix B. If outputting all occurrences of every SMEM within *T* is also desired, this is possible in *O*(*r*) space and *O*(*r* + *occ*) time with a small modification to Algorithm 1 using *ϕ* and *ϕ*^*−*1^. See Algorithm 2 for the pseudocode. These results are summarized as:

**Figure 1:**
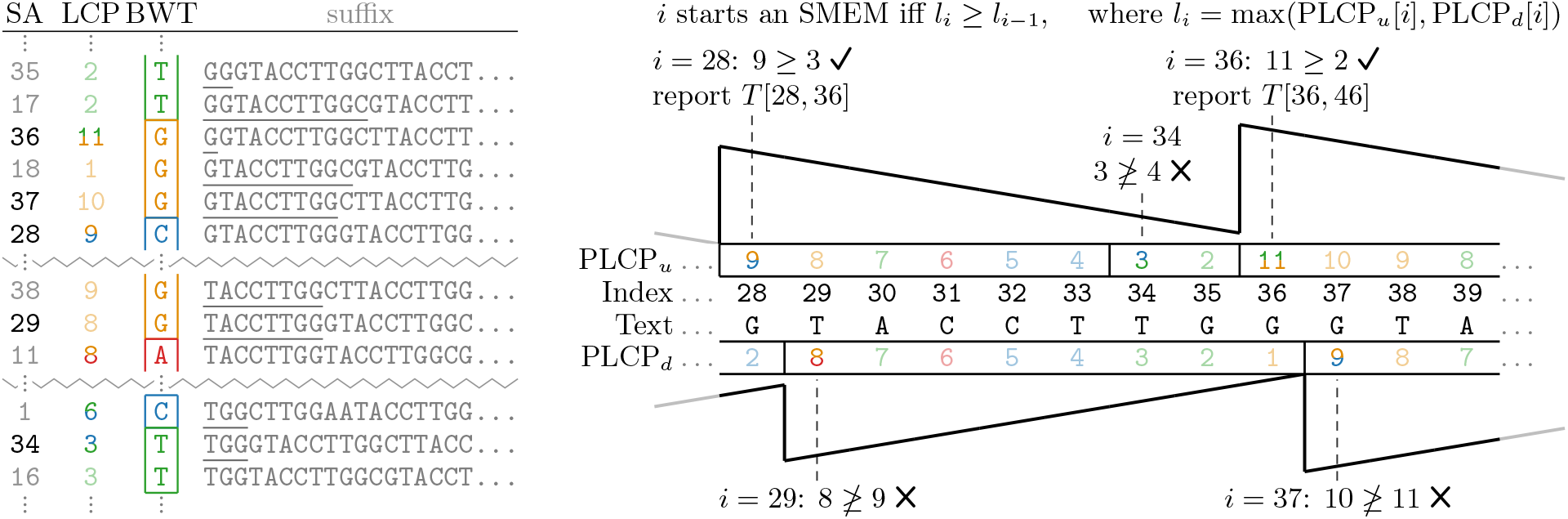
SMEM algorithm. Black values in the suffix array are those that occur in *I*^*u*^ or *I*^*d*^ (are at the top or bottom of a run) and are in [28, 39]. Here, *I*^*u*^ = {…, 28, 34, 36, … *}* and *I*^*d*^ = {…, 29, 37}, Adjacent suffixes have their *lcp*_*l*_ values (*LCP* value of the lexicographically larger suffix) drawn as lines between them. Out of 12 positions in the text, only 5 needed to be checked for starting an SMEM.

#### Theorem 1

*Algorithm 1 computes all SMEMs in a text T in O*(*r*) *space and time given an O*(*r*) *space data structure as input*.

*Proof*. The data structures required as input are *I*^*u*^, *I*^*d*^, *RP LCP*_*u*_, and *RP LCP*_*d*_, all of size *O*(*r*). The algorithm uses *O*(1) auxiliary space and *O*(1) operations per element of *I*^*u*^ and *I*^*d*^, for a total of *O*(*r*) time and space. It computes all SMEMs of *T* by Lemmas 1 and 2.

#### Theorem 2

*Algorithm 2 computes all SMEMs in a text T and all of their occurrences in O*(*r*) *space and O*(*r* + *occ*) *time given an O*(*r*) *space data structure as input*.

*Proof*. In addition to the input of Algorithm 1, Algorithm 2 only requires the *ϕ* and *ϕ*^*−*1^ move structures described in Claims 1 and 2, each *O*(*r*) space. The computations that Algorithm 2 does that Algorithm 1 doesn’t do are all in lines 5 - 13, the reporting of occurrences. These computations are overall constant per occurrence with the addition of a constant overhead per SMEM, for a total of *O*(*r* + *occ*) time for the whole algorithm. This algorithm outputs all SMEMs by Lemmas 1 and 2, and all of their occurrences by the fact that for any SMEM *T* [*i, i* + *l −* 1], all suffixes *T* [*i*^*′*^, *n −* 1] that start with *T* [*i, i* + *l −* 1] (i.e. *i*^*′*^ s.t. *lcp*_*l*_(*T* [*i*^*′*^, *n −* 1], *T* [*i, n −* 1]) *≥ l*) occur adjacent to each other in the suffix array. Lines 5 - 13 of the algorithm find all of these in constant time per occurrence by traversing up the suffix array with *ϕ* and down with *ϕ*^*−*1^.

### 3.2 Maximal Exact Matches Algorithm

We present an algorithm that outputs all MEMs of length *L* or longer in *O*(*r*) space and *O*(*r* + *occ*) time given an *RLBW T* and an *LCP*_*r*_ of the text. This algorithm utilizes the following two properties:

#### Lemma 3

*For any MEM, T* [*i, j*] = *T* [*i*^*′*^, *j*^*′*^], *let SA*[*ι*] = *i and SA*[*ι*^*′*^] = *i*^*′*^. *WLOG, ι < ι*^*′*^. *Then there exists*

*k ∈* [0, *r −* 1] *such that ι < r*_*k*_.*s ≤ ι*^*′*^.

*Proof*. If *i* 0 and *i*^*′*^ *≠* 0, then *T* [*i −* 1] *≠ T* [*i*^*′*^ *−* 1] by the definition of MEM. *T* [*i −* 1] *≠ T* [*i*^*′*^ *−* 1] implies *BWT* [*ι*] *BWT* [*ι*^*′*^] by the definition of *BWT* . If *i* = 0, *i*^*′*^ *≠* 0 by the definition of MEM and *WT* [*ι*] = 0 *≠ BWT* [*ι*^*′*^] by the fact that 0 occurs exactly once in the *BWT* . If *i*^*′*^ = 0, *i ≠* 0 by the definition of MEM and *BWT* [*ι*^*′*^] = 0*≠ BWT* [*ι*] by the fact that 0 occurs exactly once in the *BWT* . Since *BWT* [*ι*]*≠ BWT* [*ι*^*′*^], *ι* and *ι*^*′*^ must be contained in distinct runs in the *BWT* . The fact that *BWT* [*ι*] and *BWT* [*ι*^*′*^] are in distinct runs and the runs in the *BWT* are disjoint and fully cover [0, *n−*1] straightforwardly implies there exists *k ∈* [0, *r −* 1] such that *ι < r*_*k*_.*s ≤ ι*^*′*^.

#### Lemma 4

*For any MEM, T* [*i, j*] = *T* [*i*^*′*^, *j*^*′*^], *let SA*[*ι*] = *i and SA*[*ι*^*′*^] = *i*^*′*^. *WLOG, ι < ι*^*′*^. *Then* min*{LCP* [*k*]: *ι < k ≤ ι*^*′*^*}* = *j − i* + 1.

*Proof*. Let *SA*[*ρ*] = *p* and *SA*[*ρ*^*′*^] = *p*^*′*^ for any *ρ < ρ*^*′*^. Then *lcp*_*l*_(*T* [*p, n −* 1], *T* [*p*^*′*^, *n −* 1]) = min*{LCP* [*k*]: *ρ < k ≤ ρ*^*′*^*}*. Since *T* [*i, j*] = *T* [*i*^*′*^, *j*^*′*^], *lcp*_*l*_(*T* [*i, n −* 1], *T* [*i*^*′*^, *n −* 1]) = *j − i* + 1. Therefore min*{LCP* [*k*]: *ι < k ≤ ι*^*′*^*}* = *j − i* + 1.

The algorithm for outputting all MEMs of length *L* or longer processes the maximal runs of the *RLBW T* from run 0 to run *r −* 1 (ordered from smallest to largest starting position). For each run, for each suffix *d* in the run (from lexicographically smallest to largest), we output all MEMs *T* [*d*, ..] = *T* [*u*, ..] where *T* [*u, n −* 1] is lexicographically smaller than *T* [*d, n −* 1]. For some *d*, all possible suffix *u*-s are processed from lexicographically largest to smallest. The local suffix and *LCP* arrays are recovered starting from *s*, the suffix at the top of the run using the *ϕ/ϕ*^*−*1^*/P LCP*_*u*_*/P LCP*_*d*_ computing structures in *LCP*_*r*_. The minimum LCP value between the suffix at top of the run (*s*) and the current suffix above the run (*u*) is maintained as *l*_*u*_ = *lcp*_*l*_(*T* [*s, n −* 1], *T* [*u, n −* 1]). The minimum LCP value between the suffix at the top of the run (*s*) and the current suffix in the middle of the run (*d*) is also maintained as *l*_*d*_ = *lcp*_*l*_(*T* [*s, n −* 1], *T* [*d, n −* 1]). Then *lcp*_*l*_(*T* [*u, n −* 1], *T* [*d, n −* 1]) = min(*l*_*u*_, *l*_*d*_). Finally, suffixes in runs with the same RLBWT character are skipped by maintaining the minimum LCP value of each run processed so far as *r*_*k*_.*mLCP d*. This process is depicted in Figure 2. High-level pseudocode of this algorithm is provided in Algorithm 3 in Appendix B.

**Figure 2:**
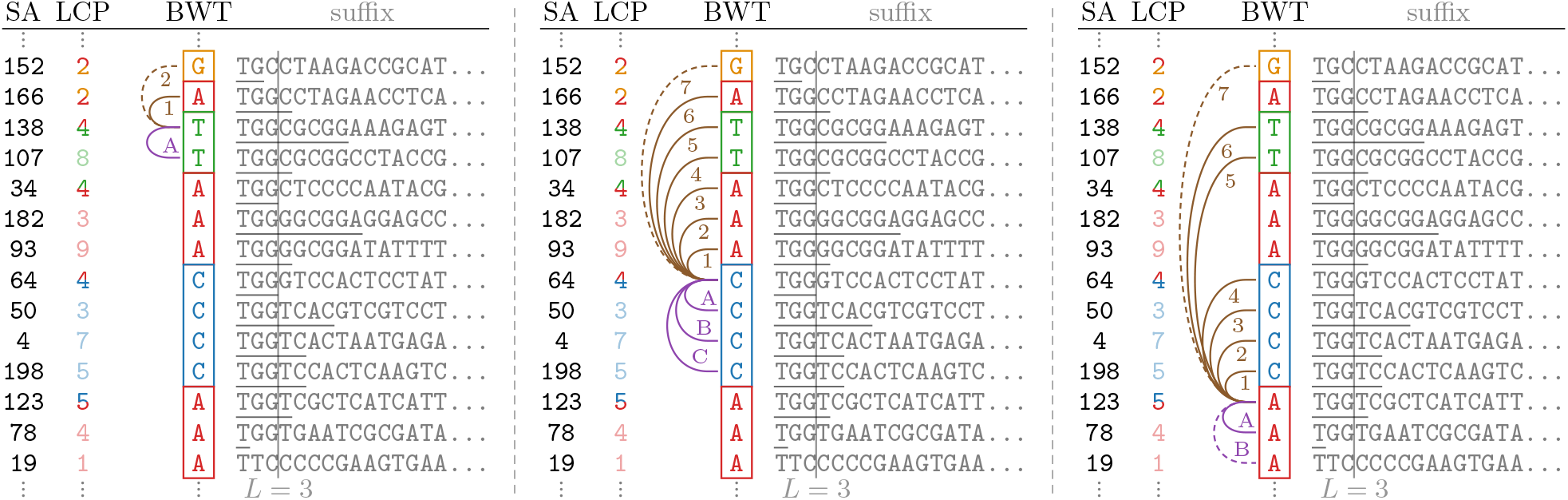
Outputting all MEMs for multiple runs in a *BWT* . For each run, the suffix at the top of the run is used as a pivot, and all *LCPs* between it and suffixes above it are computed (depicted as brown arcs) as are all *LCPs* between the top of the run and suffixes below it (purple arcs). Dashed lines are *LCP* s calculated that are lower than *L* and therefore terminate a traversal. Runs of the same value are skipped in constant time using previously computed minimum LCPs per run (see the rightmost panel). In the middle panel, the sequence of computations is 1234567A1234567B1234567C1234567.

#### Theorem 3

*Given a length threshold L and an O*(*r*) *space index of T constructed independently of L, Algorithm 3 outputs all MEMs within the text of length L or longer in O*(*r* + *occ*) *time and O*(*r*) *space where occ is the number of MEMs outputted*.

*Proof*. Algorithm 3 only requires *LCP*_*r*_ and the *RLBW T* as input, both of which are size *O*(*r*). By Lemmas 3 and 4, this algorithm outputs all MEMs of length *L* or more. The working space of the algorithm is *O*(*r*) since the input is *O*(*r*), and on top of the input, it maintains a constant number of values plus one value per run.

For all values of *d* across all runs for which there exists a MEM *T* [*d*, ..] = *T* [*u*, ..] with some lexicographically smaller suffix *u*, a constant number of operations are computed per MEM outputted, and at most one suffix *u* without a MEM is evaluated per *d*, so these operations are bounded by *O*(*occ*). Furthermore, per run at most one suffix *d* without any MEMs *T* [*d*, ..] = *T* [*u*, ..] to a lexicographically smaller suffix *u* is evaluated, and this evaluation takes a constant number of operations, so these operations are bounded by *O*(*r*). The only operations still not bounded by a constant multiple of *occ* or *r* are the operations associated with skipping runs of the same character. For any maximal run in the *BWT* to be reached by *u*, and skipped, there must have been a suffix immediately before, *ϕ*^*−*1^[*u*], for which a MEM was outputted (since *T* [*ϕ*^*−*1^[*u*] *−* 1 mod *n*] ≠ *T* [*u −* 1 mod *n*]). Therefore the number of runs skipped is also bounded by *occ* and the run time of Algorithm 3 is *O*(*r* + *occ*).

## 4 Results

We implemented the algorithms for computing MEMs and SMEMs and their occurrences in *O*(*r*) space and *O*(*r* + *occ*) time as `TeraMEM` in the TeraTools suite available at https://github.com/ucfcbb/TeraTools. In Section 4.1, we benchmark runtime and memory usage for computing MEMs across all pairs of sequences in large repetitive datasets. In Section 4.2, we highlight potential applications of all-vs-all MEM analyses by presenting discoveries found in a well-assembled human centromere dataset.

We use three datasets in our computational experiments: `mtb152, humCent625`, and `openHGL. mtb152` is a dataset of 152 Mycobacterium tuberculosis complete genome assemblies, 151 of which were assembled by Marin et al. [25], and the last (H37Rv) is a reference assembly by Cole et al. [9]. It is available at https://doi.org/10.5281/zenodo.18946124. humCent625 is a set of human centromere assemblies from 625 haplotypes aggregated by Gao et al. [16], downloadable from https://doi.org/10.6084/m9.figshare.29921990. humCent625 is composed of 5,747 centromeres from 453 haplotypes from the Human Pangenome Reference Consortium (HPRC) release 2 dataset [24], 2,104 centromeres from 130 haplotypes from the Human Genome Structural Variation Consortium phase 3 dataset [23], and 396 centromeres from 42 haplotypes from the Platinum Pedigree Consortium dataset [30]. Finally, openHGL is a dataset of 579 high-quality human assemblies consisting of the HPRC release 2 [24] dataset, the Arab Pangenome Reference [26], and multiple reference genomes [32, 40, 20, 35, 4, 8]. It is available at https://doi.org/10.5281/zenodo.11533210.

For each dataset, a text is generated by concatenating all strings and their reverse complements separated by string termination characters. Then, all Ns are removed and split contigs. Statistics for these datasets can be seen in Table 1.

**Table 1:** Statistics for tested datasets.

|  | *n* | *h* | *N* | *r* | $\frac{n}{r}$ |
|---|---|---|---|---|---|
| mtb152 | 1,341,467,444 | 152 | 304 | 6,740,684 | 199.0 |
| humCent625 | 72,636,274,842 | 625 | 16,494 | 166,446,671 | 436.4 |
| openHGL | 3,483,676,117,190 | 579 | 117,392 | 5,507,663,548 | 632.5 |

*n*: length of text (number of basepairs and string termination characters). *h*: number of haplotypes. *N*: number of strings, forward and reverse complements counted separately. *r*: runs in the *BWT*. $\frac{n}{r}$: average run length in the *BWT*.

### 4.1 Run time and Memory

We evaluate three methods for computing all-vs-all pairwise matches: `copMEM2` [17], `Mumemto` [37], and `TeraMEM` (this paper). `Mumemto` is the only tool we found computing all-vs-all matching information in assemblies in linear time and compressed space. `copMEM2` is the fastest and most recent tool we found for computing MEMs between a pair of genomes. `Mumemto` and `TeraMEM` compute all-vs-all MEMs natively and both utilize a precomputed compressed index as input. `Mumemto` takes as input a prefix-free parse (*PFP*) [5] of the text while `TeraMEM` takes as input an *RLBW T* -based index of the text. It has been shown that |*PFP*| can be 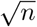 times larger than *r* [27, 18]. We simulate an all-vs-all method based on `copMEM2` simply by calling it on all pairs of genomes in a dataset. Finally, note that we compare to `Mumemto’s` partial multi-MEM outputting mode. A partial multi-MEM is: a string *T* [*i, j*] that occurs in at least *k* distinct genomes and there exists a MEM *T* [*i, j*] = *T* [*i*^*′*^, *j*^*′*^] across at least two genomes. We set *k* = 2 and a minimum length of 1 million basepairs for all matches to minimize differences between partial multi-MEMs and pairwise MEMs.

All experiments are performed on a server with an Intel(R) Xeon(R) Gold 6338N CPU with 128 cores and 1 TiB of RAM. The times and memory usages reported here are the elapsed wall clock times and maximum resident set sizes reported by `/usr/bin/time -v`. Since `Mumemto` is sequential and `TeraMEM` is multi-threaded, we evaluate `TeraMEM` and `copMEM2` as both single-threaded and with all available cores for all experiments.

The run time for computing MEMs from the text is provided in Table 2 and the peak memory is provided in Table 3. The run time is divided into two parts, a one-time index construction cost for `TeraMEM` and `Mumemto`, and the MEM computing cost given the required input for all tools. Some values are estimated for costly experiments. A full breakdown of the estimation methods is provided in Appendix C. The `TeraMEM` pipeline for computing MEMs from the text is lg *→* TeraLCP *→* TeraIndex *→* TeraMEM. lg (lyndongram-mar [29]) constructs the RLBWT from the text. `TeraLCP` and `TeraIndex` construct the RLBWT-based compressed index that `TeraMEM` takes as input and are from TeraTools [34]. Compared to `Mumemto`, `TeraMEM` uses 7.5 times less memory and is 14.4% slower single-threaded but 20 times faster given multiple cores on `humCent625`. Compared to copMEM2, `TeraMEM` is up to 27 times more memory efficient and 16 times faster on humCent625. `TeraMEM’s` run time scales very linearly by number of cores, so 2 cores is enough to beat `Mumemto’s` run time in both mtb152 and `humCent625`. On `openHGL`, `copMEM2` uses up to 92% more memory but is around 70% faster than the full `TeraMEM` pipeline when computing MEMs from the text directly. We address this below.

**Table 2:** Run time: index construction + computing MEMs.

| tool | cores | mtb152 | humCent625 | openHGL |
|---|---|---|---|---|
| copMEM2 | 1 | 0.00 + 13,781.00 | 0 + 278,396<sup>†</sup> | - |
|  | 128 | 0.00 + 846.49 | 0 + 16,277 | 0 + 52,382<sup>†</sup> |
| *PFP* + Mumemto | 1 | 51.78 + 214.45 | 2,980 + 17,690 | ≥132,379<sup>†</sup> + ≥556,000<sup>†</sup> |
| *LCP<sub>r</sub>* + TeraMEM | 1 | 274.84 + 5.11 | 23,598 + 56 | - + 6,972 |
|  | 128 | 24.99 + 0.25 | 1,007 + 7 | 90,828 + 279 |

Reported times are in seconds. <sup>†</sup>: estimated value, see Appendix C. The *LCP<sub>r</sub>* is constructed by lg → TeraLCP → TeraIndex.

**Table 3:** Peak Memory during index construction and MEM computation.

| tool | cores | mtb152 | humCent625 | openHGL |
|---|---|---|---|---|
| copMEM2 | 1 | 2.04 | 2.15<sup>†</sup> | - |
|  | 128 | ≤261.53<sup>†</sup> | ≤274.66<sup>†</sup> | ≤659<sup>†</sup> |
| *PFP* + Mumemto | 1 | 1.06 | 76.45 | ≥2,765<sup>†</sup> |
| *LCP<sub>r</sub>* + TeraMEM | 1 | 0.33 | 9.42 | 343<sup>†</sup> |
|  | 128 | 0.66 | 10.13 | 343 |

Reported in GiB. <sup>†</sup>: estimated value, see Appendix C.

#### TeraMEM vs. copMEM2 on Pangenomes

The vast majority of the time cost for `TeraMEM` occurs in the linear time compressed index construction step. As this is a one-time cost, if MEMs are to be called repeatedly from the dataset, `TeraMEM` is many times more efficient than `copMEM2`. A plausible scenario where repeatedly calling MEMs is useful is when the number of MEMs is much larger than the size of the compressed index, this occurs when the length threshold is small. This is useful when filtering MEMs down to those with properties that occur rarely (the utility of this is demonstrated in Section 4.2). Furthermore, since the runtime of `copMEM2` scales quadratically as opposed to the linear time cost of index construction, the run times of these tools equilibrate at 1,000 haplotypes: copMEM2’s run time would increase 3 times, to 157k seconds, while `TeraMEM’s` runtime would increase 72.7%, also to 157k seconds. Past that size, the index construction of `TeraMEM` will be cheaper than computing MEMs with `copMEM2`. Finally, the benchmark in Table 2 is overly generous to `copMEM2`, by only calling MEMs between pairs of genomes, it fails to output MEMs between contigs of the same haplotype and MEMs within a contig (`Mumemto` also fails to output these). The benefit of computing these types of matches is shown in Section 4.2. For `copMEM2` to output the same MEMs `TeraMEM` outputs, it must be called on all pairs of contigs. This slows copMEM2 down thousands of times. It took more than two hours to compute MEMs in a dataset of 5 haplotypes with 128 cores, implying it would take over a thousand days to correctly output all MEMs in `openHGL`.

#### Run time and Memory Scaling

We measure the scaling of run time and memory usage as datasets grow larger and more repetitive. This is done by randomly downsampling mtb152 and humCent625. The downsampling is done by generating a random order of the haplotypes in each dataset. Then, the first 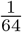 of the haplotypes form the smallest downsampling, the first 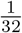 form the second smallest downsampling, and so on 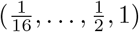. Therefore each downsampled dataset is a subset of all larger downsampled datasets. In Figure 3, we plot the % increase in run time and memory requirements relative to a dataset half the size for computing MEMs from a compressed index for subsets of `humCent625`. In a similar fashion, Figure A9 plots the scaling of resource requirements for computing MEMs directly from the text. The run time of Mumemto scales linearly in both settings, however the run time of `TeraMEM` scales clearly sublinearly when computing MEMs from a compressed index. When computing MEMs from the text, `TeraMEM` scales linearly. It seems like the run time of parallelized `TeraMEM` scales sublinearly, but this is likely caused by the runtime being dominated by fixed cost operations since the parallelized pipeline is so fast. Since the parallelized `TeraMEM` finishes in a few seconds, a significant portion of the time is file I/O. The memory usage of `Mumemto` and `TeraMEM` both scale sublinearly, but `TeraMEM’s` memory usage grows much slower than `Mumemto’s`. These effects compound over many doublings of the dataset (after 6 doublings, Mumemto’s memory usage was 17.2 times larger than originally while `TeraMEM’s` was 5.4 times larger) and are expected to continue as datasets grow past current sizes. Plots for the run time, memory usage, and their scaling for `mtb152` and `humCent625` for computing MEMs directly from the text or from a compressed index can be seen in Appendix D.

**Figure 3:**
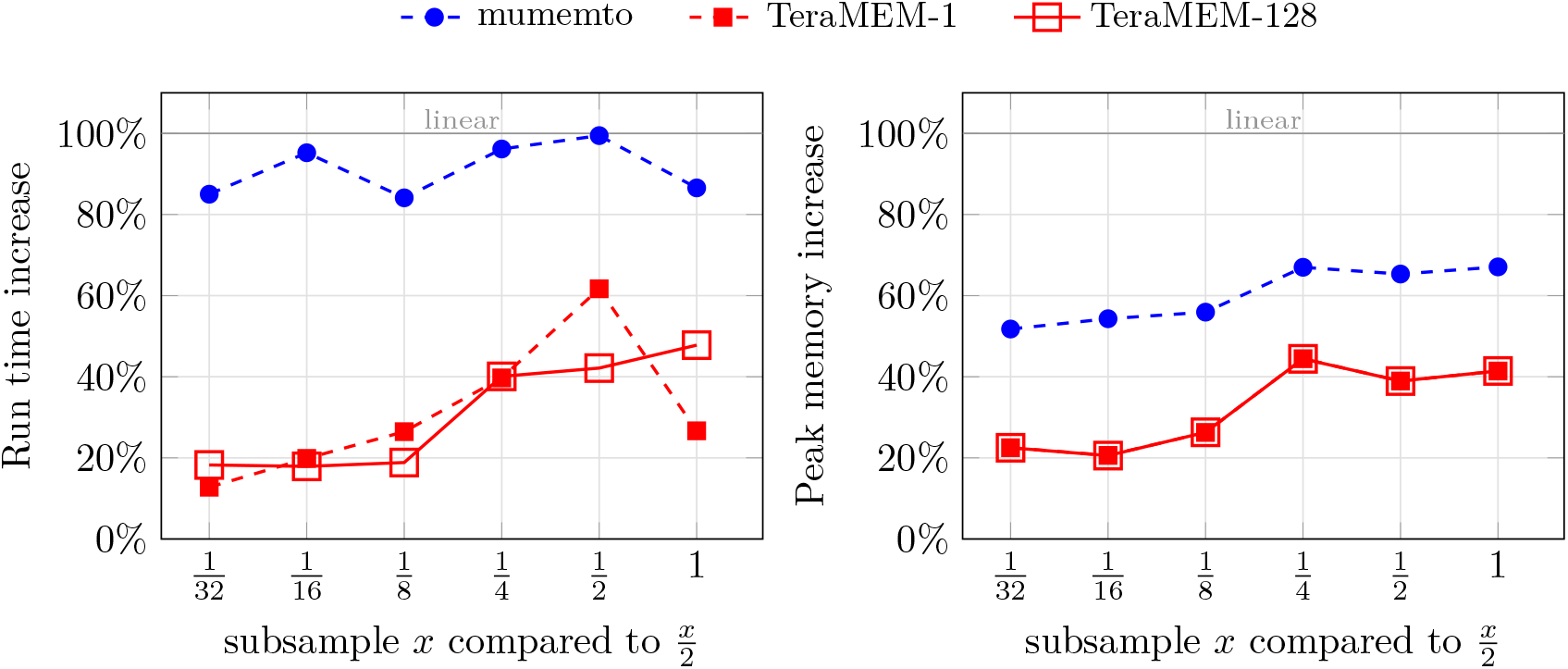
Resource requirement scaling for computing MEMs from a compressed index as datasets double in size for subsets of humCent625. Time on the left and memory on the right. X-axis is size of the twice as large dataset, y-axis is the % additional resource required relative to the subsampled dataset half the size. 100% additional resources is linear growth, *<* 100% is sublinear.

### 4.2 MEMs in Human Centromeres

We attempted to use all-vs-all pairwise MEMs to find “interesting” pairs of centromeres. We do this by first calling all-vs-all MEMs of length at least 10 kbp among all pairs of centromeres in the HPRC release 2 dataset [24] annotated by and provided in Gao et al. [16]. We filter these MEMs down to those exhibiting any of the following four properties:

- Interchromosomal: MEMs between centromeres of different chromosomes
- Inverted: MEMs between opposite strand orientations: one forward strand and one reverse strand
- Self: MEMs within the same contig (duplication)
- Self inverted: MEMs within the same contig on opposite strands

We plot the MEMs for a pair of contigs using SVbyEye [31]. For interchromosomal and inverted MEMs, we plot the 5 pairs of centromeres with the greatest total length of MEMs with each property for every pair of chromosomes. Similarly for self and self inverted MEMs: for every chromosome, we plot the 5 centromeres with the largest total length of MEMs with each property. These plots are then visually examined. In providing examples of the detected phenomena, we add dot plots by ModDotPlot [39] to confirm MEM-identified structures and provide finer-scale details.

We find multiple Chromosome 13 and 21 centromeres with MEMs 10 kbp or larger spanning the centromeres, both in and out of the active *α*-satellite higher-order repeat (HOR) arrays. This phenomenon was recently discovered by Eizenga et al. [14] and is possibly caused by recent interchromosomal exchange of whole centromeres. Eizenga et al. also found relationships between the centromeres of Chromosomes 14 and 22. Chromosomes 14 and 22 were the only other pair of chromosomes with interchromosomal MEMs larger than 10 kbp, however the pairs with the most total matching length only had a few matches totalling a few tens of kbp, possibly indicating our MEM length threshold is too large or the 14–22 relationship too old to detect with long exact matches directly.

Chromosomes 4, 17, and X have inverted MEMs but don’t have strong visual support of an inversion. These inverted MEMs are either isolated, occur on less than 5 pairs of centromeres, or occur in inverted repeats. On the other hand, we find sequences of inverted MEMs that do seem to correspond to inversions for Chromosomes 1, 2, 7, 11, 14, 20, and 22. These inversions are typically at least 100 kbp. Inverted MEMs on Chromosome 1 reveal a known [1] *>*1 Mbp inversion within the active *α*-satellite HOR array. Inverted MEMs on Chromosomes 2, 7, 11, 14, 20, and 22 all occur just outside the active *α*-satellite HOR array. The Chromosome 2 active *α*-satellite HOR array is flanked by a *∼*200–300 kbp inversion on one side and a *∼* 200–300 kbp pseudopalindrome (*∼* 10 kbp hsat2 spacer surrounded by a *∼* 120 kbp inverted repeat, see Figure 4a). Chromosomes 7, 11, 14, 20, and 22 each have one inversion of a hundred to a few hundred kbp outside of the active *α*-satellite HOR array.

**Figure 4:**
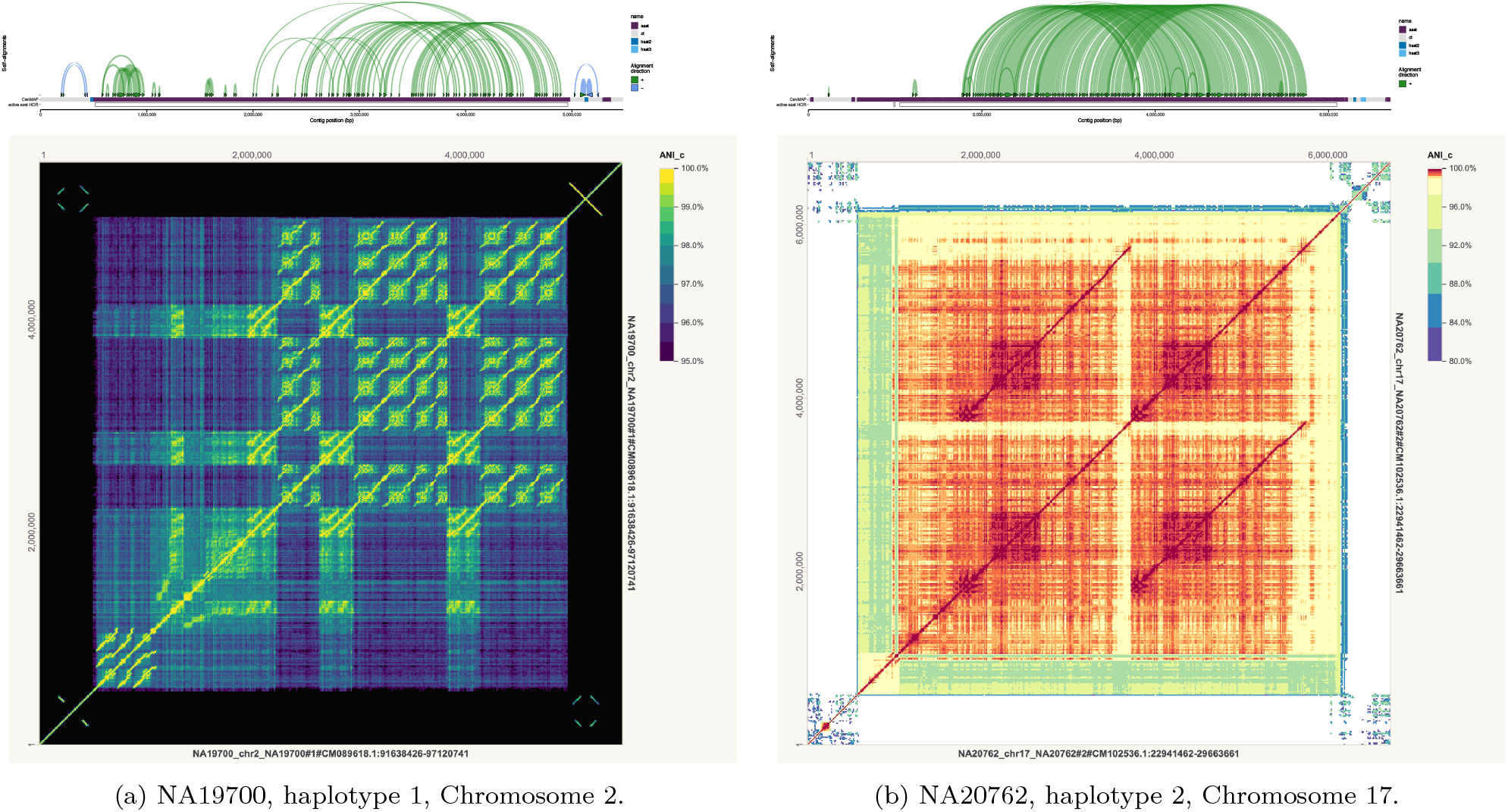
Segmental duplications in Chromosomes 2 and 17. The pseudopalindrome and inverted repeat flanking the Chromosome 2 active *α*-satellite HOR array are also visible. The top plot is the SVbyEye [31] plot where each block is a MEM of length at least 10 kbp. The bottom plot is a dot plot generated by ModDotPlot [39].

When plotting the 5 centromeres per chromosome with the longest total length of MEMs with themselves on the same contig and strand, we observe long tracts of self-MEMs that seem to indicate large segmental duplications. We find these segmental duplications in every chromosome except Chromosome Y. They range from a few hundred thousand to millions of basepairs in length and occur both inside and outside of the active *α*-satellite HOR arrays, see Figure 4.

In plotting inverted self MEMs, we find inverted repeats and pseudopalindromes in or around the centromeres of Chromosomes 2, 4, 7, 11, 20, 22, and X. Chromosomes 2 and 7 have a pseudopalindrome on one side and an inverted repeat on the other. Chromosomes 4 and X have a pseudopalindrome on one side. Chromosomes 11, 20, and 22 have one small inverted repeat. Large pseudopalindromes in the sex chromosomes are well established [19, 32], but may not be in autosomal pericentromeric regions. See Appendix E for additional plots.

## 5 Discussion

In this work, we have developed algorithms for outputting all Maximal Exact Matches (MEMs) or Super-maximal Exact Matches (SMEMs) and their occurrences in time proportional to a dataset’s compressed size and the number of outputted matches given a compressed index as input. These algorithms run thousands of times faster and use less than one seventh the memory of previous compressed methods and fastest pair-wise algorithms on current pangenomes. As pangenomes grow from their current hundreds of haplotypes to planned thousands of haplotypes in the next couple of years, the gap in run time and memory usage between the algorithms presented here and previous best algorithms will grow as well. In addition, we have shown the utility of pairwise MEM-based analyses through an application to centromeres of a few hundred human haplotypes. This analysis revealed previously unknown structures within and around centromeres: large segmental duplications, inverted repeats, and pseudopalindromes. Therefore, MEM-based analyses may be fruitful in other contexts as well, including genome-wide analyses or analyses in other species.

There are multiple avenues for possible future work. On the computational side, the general technique for computing MEMs and SMEMs in sublinear time and compressed space developed here is likely extendible to other useful classes of matches, for example multi-MEMs, multi-MUMs, partial multi-MEMs/MUMs [37], and others. These techniques also apply to other structures used in bioinformatics related to the Burrows-Wheeler transform: the positional Burrows-Wheeler transform [12] and the Graph Burrows-Wheeler transform [38]; future work could develop these sublinear time and compressed space matching algorithms on these structures as well. Furthermore, MEM algorithms based on non-RLBWT-based compression schemes are unimplemented and underexplored; for example, the grammar-based MEM algorithm by Díaz-Domínguez and Salmela [10] might outperform ours, but is unimplemented. Finally, RLBWT-based methods induce one random access per character searched or MEM outputted, implying a fundamental speed limit: RAM throughput. Cache-efficient compressed indexes [2] or techniques for combining multiple bases into one BWT character, such as syncmers or pangenome graphs [13, 11, 38] may overcome this limitation.

On the biological side, software for conducting MEM-analyses such as the one we conducted by eye in Section 4.2 in a more principled, automated, and efficient fashion would be valuable. MEM analysis on other datasets, such as whole human genomes, bacterial datasets, or other species with large well-assembled pangenomes may also prove insightful. Finally, the original motivation for finding all-vs-all long exact matches in large haplotype panels was to identify long genomic segments between pairs of individuals that are identical-by-descent (IBD). Tools could be developed for identifying IBD segments directly in assemblies, in a position-agnostic and reference-unbiased manner. Indeed, this was one of the original motivations the authors had to pursue this work.

## Acknowledgments

This work was supported by the National Institutes of Health under grants R01HG010086 and U01AG098387.

## Appendices

### A Proofs

#### Claim 1

*Let* Φ(*i*) = |{*j ≤ i*: *SA*[*r*_*k*_.*s*] = *j* for some *k ∈* [0, *r −*1] *}*| *−* 1. *There exists an O*(*r*) *space data structure that computes ϕ*[*i*], Φ(*ϕ*[*i*]), *and P LCP*_*u*_[*i*] *in constant time given* Φ(*i*).

*Proof*. Recall that *ϕ* is a permutation of [0, *n −* 1]. Furthermore, for any suffix *j* = *SA*[*i*], if *i* is not the top of a maximal run in the *BWT* (i.e. if *i* ≠ *r*_*k*_.*s* for all *k*), then *ϕ*[*j −* 1] = *ϕ*[*j*] *−* 1 (see Lemma 3.5 of [15] or Lemma 3 of [33] for a proof of this property). For the same conditions on *i* and *j, P LCP*_*u*_[*j −* 1] = *P LCP*_*u*_[*j*] + 1 (see Lemma 2 in [33] for a proof of this property). Therefore, we construct a sequence of disjoint conserved intervals in *ϕ* fully covering [0, *n −* 1] by setting the starting points of the disjoint intervals to be exactly the starting points of the suffixes occurring at the tops of maximal runs, *I*^*u*^ = *SA*[*r*_*k*_.*s*]: 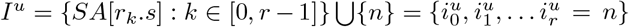. The move structure constructed on *I*^*u*^ allows the computation of *ϕ*[*i*] and Φ(*ϕ*[*i*]) in constant time and *O*(*r*) space given Φ(*i*). In addition to this move structure, we store *I*^*u*^ and *RP LCP*_*u*_, an array of length *r* where 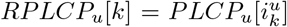. Then 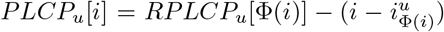 can be computed in constant time and *O*(*r*) space as well.

#### Claim 2

*Let* Φ_*inv*_(*i*) = |*{j ≤ i*: *SA*[*r*_*k*_.*e*] = *j* for some *k ∈* [0, *r −* 1] *}*|*−* 1. *There exists an O*(*r*) *space data structure that computes ϕ*^*−*1^[*i*], Φ_*inv*_(*ϕ*^*−*1^[*i*]), *and P LCP*_*d*_[*i*] *in constant time given* Φ_*inv*_(*i*).

*Proof*. The proof is the same as that of Claim 1, except replace all occurrences of *ϕ*, “top”, *I*^*u*^, 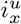, *r*_*k*_.*s, P LCP*_*u*_, Φ, and *RP LCP*_*u*_ by *ϕ*^*−*1^, “bottom”, *I*^*d*^, 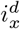, *r*_*k*_.*e, P LCP*_*d*_, Φ_*inv*_, and *RP LCP*_*d*_ respectively. See Lemma 3.5 of [15] or Lemma 4 of [28] for a proof of the corresponding *ϕ*^*−*1^ property. See Lemma 6 in [33] for a proof of the corresponding *P LCP*_*d*_ property.

#### Lemma 2

*T* [*i, j*] *is an SMEM if and only if* max(*P LCP*_*u*_[*i−*1], *P LCP*_*d*_[*i−*1]) *≤* max(*P LCP*_*u*_[*i*], *P LCP*_*d*_[*i*]) and *j* = *i* + max(*P LCP*_*u*_[*i*], *P LCP*_*d*_[*i*]) *−* 1.

*Proof*. We first show max(*P LCP*_*u*_[*i −* 1], *P LCP*_*d*_[*i −* 1]) *≤* max(*P LCP*_*u*_[*i*], *PLCP*_*d*_[*i*]) implies *T* [*i, j* = *i* + max(*P LCP*_*u*_[*i*], *P LCP*_*d*_[*i*]) *−* 1] is an SMEM. max(*P LCP*_*u*_[*i*], *P LCP*_*d*_[*i*]) = *j − i* + 1 implies that *occ*(*T* [*i, j*], *T*) *>* 1 since the length of the longest common prefix of *T* [*i, n −* 1] and the suffix above or below it in the suffix array is *j − i* + 1 = |*T* [*i, j*]|. Furthermore, max(*P LCP*_*u*_[*i*], *PLCP*_*d*_[*i*]) = *j − i* + 1 implies *occ*(*T* [*i, j* + 1], *T*) = 1 since if *T* [*i, j* + 1] occurred in *T* at the beginning of some suffix other than *T* [*i, n −* 1], that suffix would sort adjacent to *T* [*i, n −* 1] in the suffix array and either the *P LCP*_*u*_[*i*] or *P LCP*_*d*_[*i*] value would be at least *j − i* + 2, the length of *T* [*i, j* + 1]. Finally, max(*P LCP*_*u*_[*i −* 1], *P LCP*_*d*_[*i −* 1]) *≤* max(*P LCP*_*u*_[*i*], *PLCP*_*d*_[*i*]) implies *occ*(*T* [*i −* 1, *j*], *T*) = 1 since if there existed a suffix of *T* other than *T* [*i −* 1, *n −* 1] prefixed by *T* [*i −* 1, *j*], then it would sort adjacent to *T* [*i −* 1, *n −* 1] in the suffix array and either *P LCP*_*u*_[*i −* 1] or *P LCP*_*d*_[*i −* 1] would be at least |*T* [*i −* 1, *j*] | = *j − i*+2 = max(*P LCP*_*u*_[*i*], *P LCP*_*d*_[*i*])+1 and max(*P LCP*_*u*_[*i*], *P LCP*_*d*_[*i*]) + 1 *>* max(*P LCP*_*u*_[*i*], *PLCP*_*d*_[*i*]).

Now, we show that if *T* [*i, j*] is an SMEM, max(*P LCP*_*u*_[*i−*1], *P LCP*_*d*_[*i−*1]) *≤* max(*P LCP*_*u*_[*i*], *P LCP*_*d*_[*i*]) and *j* = *i* + max(*P LCP*_*u*_[*i*], *P LCP*_*d*_[*i*]) *−* 1. We first show that *T* [*i, j*] is an SMEM ⇒ *j* = *i* + max(*P LCP*_*u*_[*i*], *PLCP*_*d*_[*i*]) *−* 1. By the fact that *occ*(*T* [*i, j*], *T*) *>* 1 and *occ*(*T* [*i, j* + 1], *T*) = 1, there exists a suffix of *T*, call it *s*, other than *T* [*i, n −* 1] such that *lcp*_*l*_(*s, T* [*i, n −* 1]) = |*T* [*i, j*]| and there does not exist any suffix *s* other than *T* [*i, n −* 1] such that *lcp*_*l*_(*s, T* [*i, n −* 1]) *>* |*T* [*i, j*]|. Therefore, by the fact that a suffix with the longest common prefix to *T* [*i, n −* 1] sorts adjacent to it in the suffix array, *j* = *i* + max(*P LCP*_*u*_[*i*], *P LCP*_*d*_[*i*]) *−* 1.

Finally, we show that max(*P LCP*_*u*_[*i −* 1], *P LCP*_*d*_[*i −* 1]) *≤* max(*P LCP*_*u*_[*i*], *PLCP*_*d*_[*i*]) is implied by *T* [*i, j*] being an SMEM. Recall that *T* [*i, j*] being an SMEM implies *occ*(*T* [*i −* 1, *j*], *T*) = 1. Therefore, max(*P LCP*_*u*_[*i −* 1], *P LCP*_*d*_[*i −* 1]) *<* |*T* [*i −* 1, *j*]| = *j − i* + 2 = max(*P LCP*_*u*_[*i*], *P LCP*_*d*_[*i*]) + 1. Therefore, max(*P LCP*_*u*_[*i −* 1], *P LCP*_*d*_[*i −* 1]) *≤* max(*P LCP*_*u*_[*i*], *P LCP*_*d*_[*i*]).

### B Pseudocode

#### Algorithm 1: SMEM algorithm

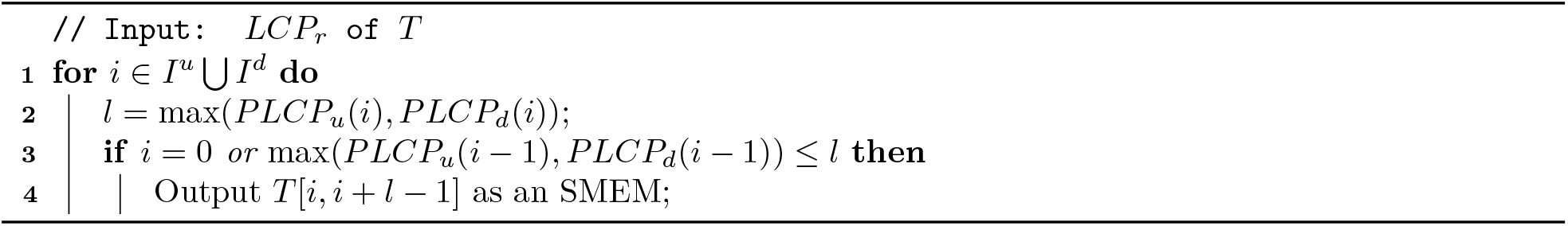

#### Algorithm 2: SMEMs and occurrences algorithm

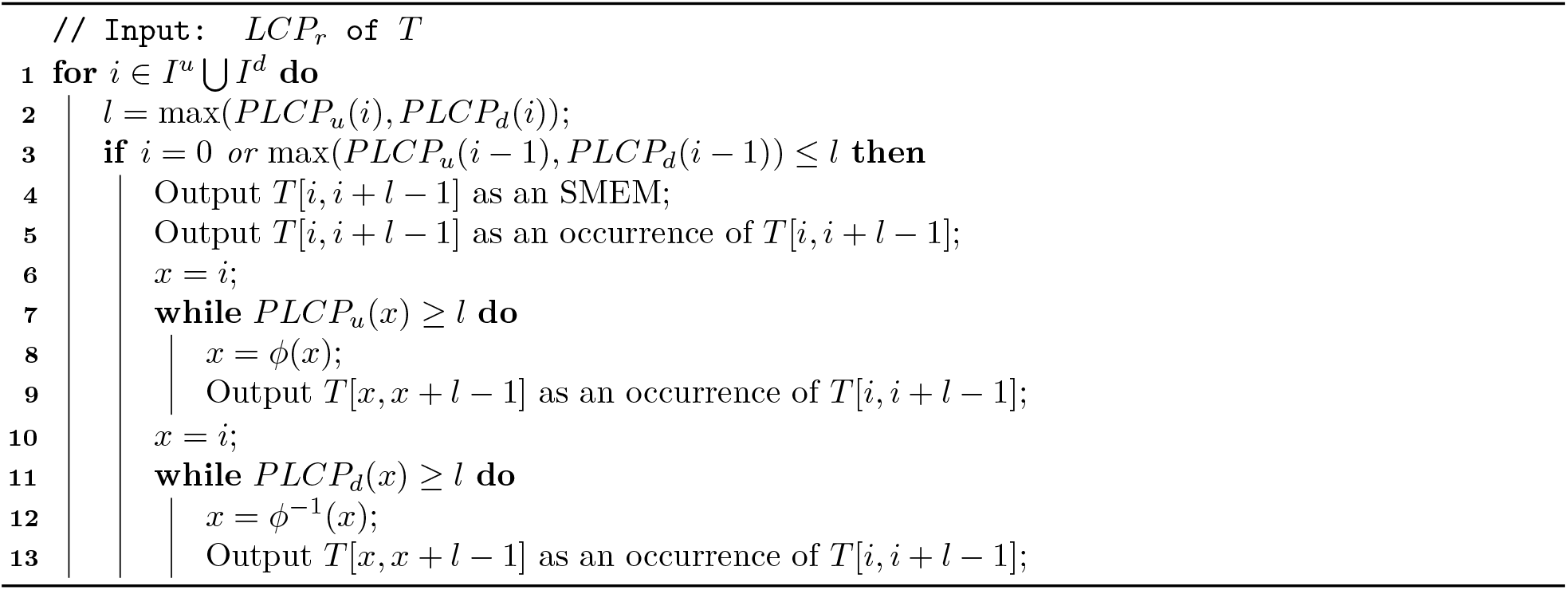

#### Algorithm 3: MEMs length *L* or longer algorithm

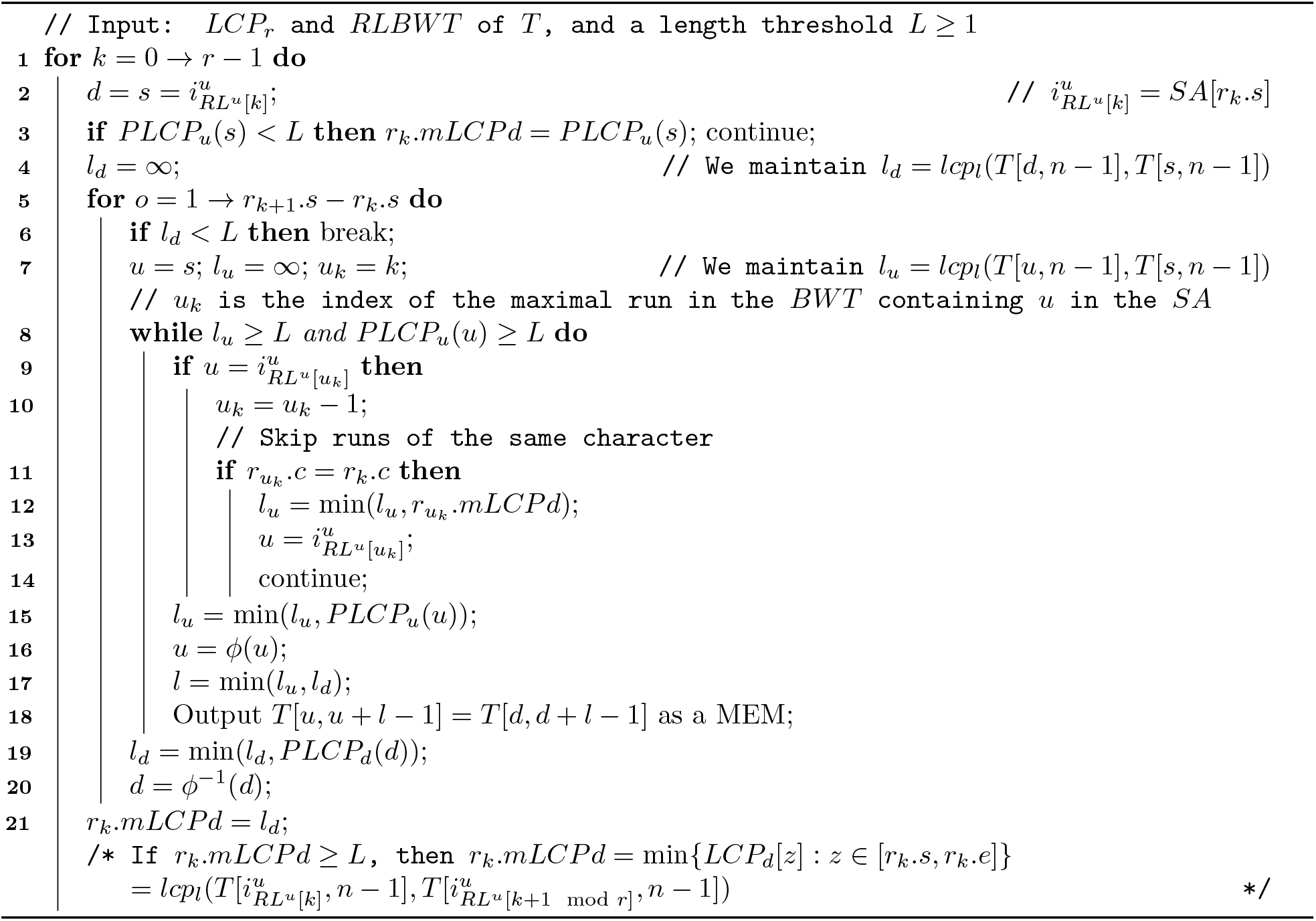

### C Benchmark Estimates

Throughout the main paper and all appendices, estimated results only occur in Tables. All figures only plot exactly measured values. Every estimate in a table is marked by a ^*†*^. Upper bounds are preceded by a less than or equal to sign and lower bounds are preceded by a greater than or equal to sign. An explanation of how every estimate was arrived to is below.

- `Mumemto`
  –Table 2: Time for computing *PFP* from the text of `openHGL. Mumemto` takes *∼* 38 ns per character on `mtb152` and *∼* 41 ns per character on `humCent625`. We use 38 ns per character as a lower bound for `openHGL`.
  –Table 2: Time for computing MEMs from a *PFP* of openHGL. `Mumemto` takes *∼* 160 ns per character on `mtb152` and *∼* 244 ns per character on `humCent625`. We use 160 ns per character as a lower bound for `openHGL`.
  –Table 3: Memory on `openHGL`. In the Results: “Anchor-based merging across 474 human haplotypes” section of [36], the authors of `Mumemto` estimate its memory usage for computing multi-maximal unique matches (multi-MUMs) on the HPRC release 2 [24] dataset. This dataset is a subset of `openHGL`, so we use their estimate as a lower bound for `Mumemto’s` memory when computing partial multi-MEMs on `openHGL`.
- `copMEM2`: For scripts spawning multiple child processes, `/usr/bin/time -v` reports the maximum memory usage of every child process of a process and the process itself but not the peak total memory usage. The peak memory values reported by `/usr/bin/time -v` for parallel and sequential calls were always equal or differed by at most 4 KiB (1 page), therefore, we treat them as equal.
  –`mtb152`, 128 cores memory: upper-bounded by 128 times the peak memory `/usr/bin/time -v` reported.
  –`humCent625`, 1 core: the dataset was subsampled to half by selecting 313 random haplotypes. Sequential pairwise `copMEM2` was run on this dataset. The estimated peak memory is the peak memory on the 313 haplotype dataset. The estimated run time is 4 times the run time on the 313 haplotype dataset.
  –`humCent625`, 128 core memory: upper-bounded by 128 times the peak memory reported by `/usr/bin/time -v`
  –`openHGL`, 128 core time: we run pairwise `copMEM2` on 36 haplotypes and multiply the time by 256.
  –`openHGL`, 128 core memory: we multiply the peak memory on the 36 haplotype sample by 128 for an upper bound.
- `TeraMEM`
  –`openHGL`, 1 core memory: estimated at the 128 core memory usage of `TeraMEM` since `TeraMEM’s` 1 core memory usage was smaller than its 128 core memory usage for `mtb152` and `humCent625`.

### D Additional Benchmarks

#### D.1 mtb152

**Figure A1:**
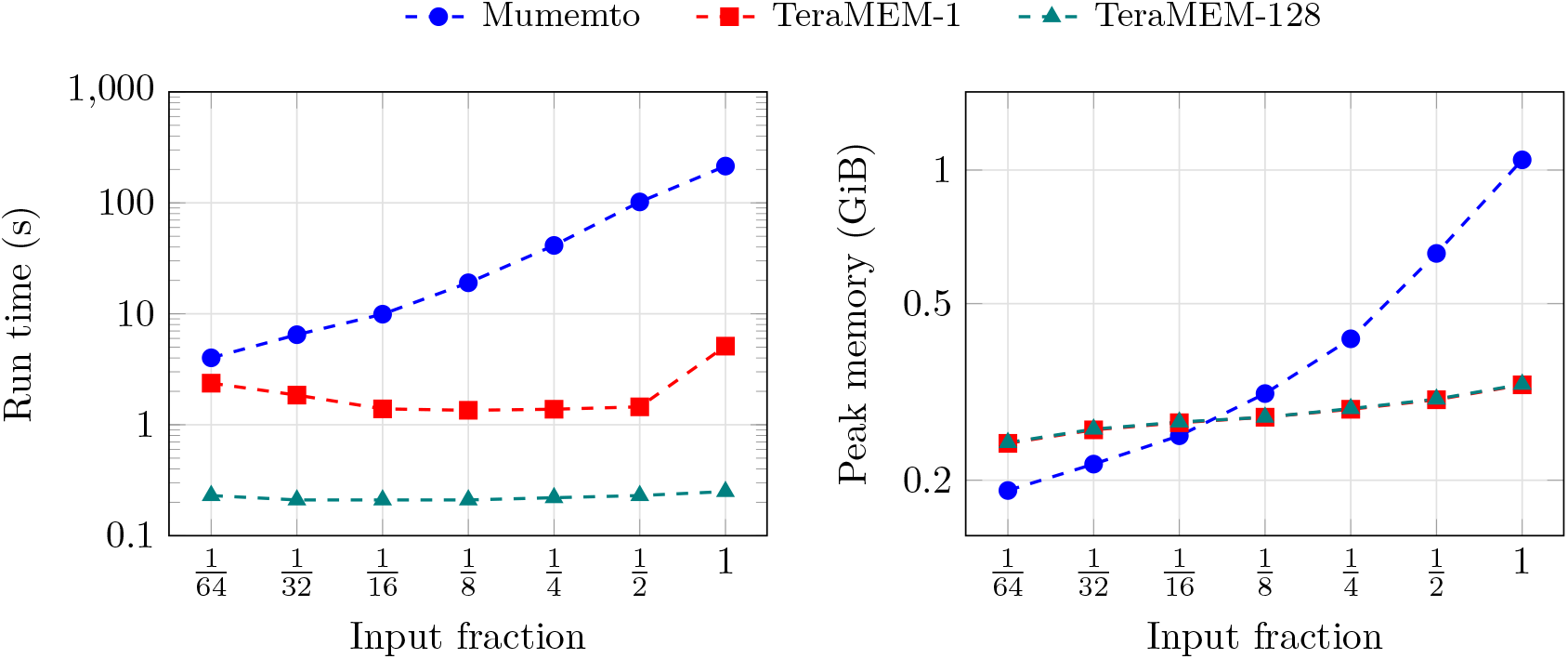
Run time and peak memory for computing MEMs from an index (`mtb152`).

**Figure A2:**
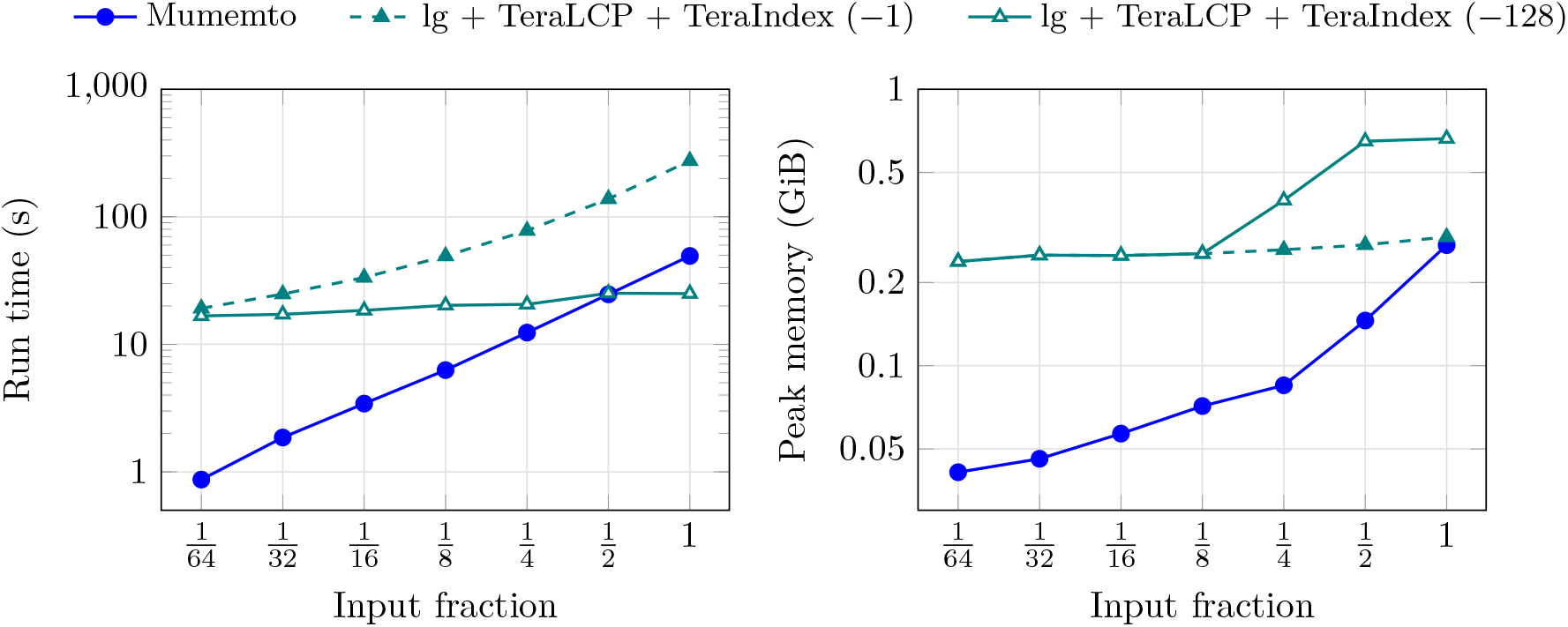
Run time and peak memory for building an index from text (`mtb152`).

**Figure A3:**
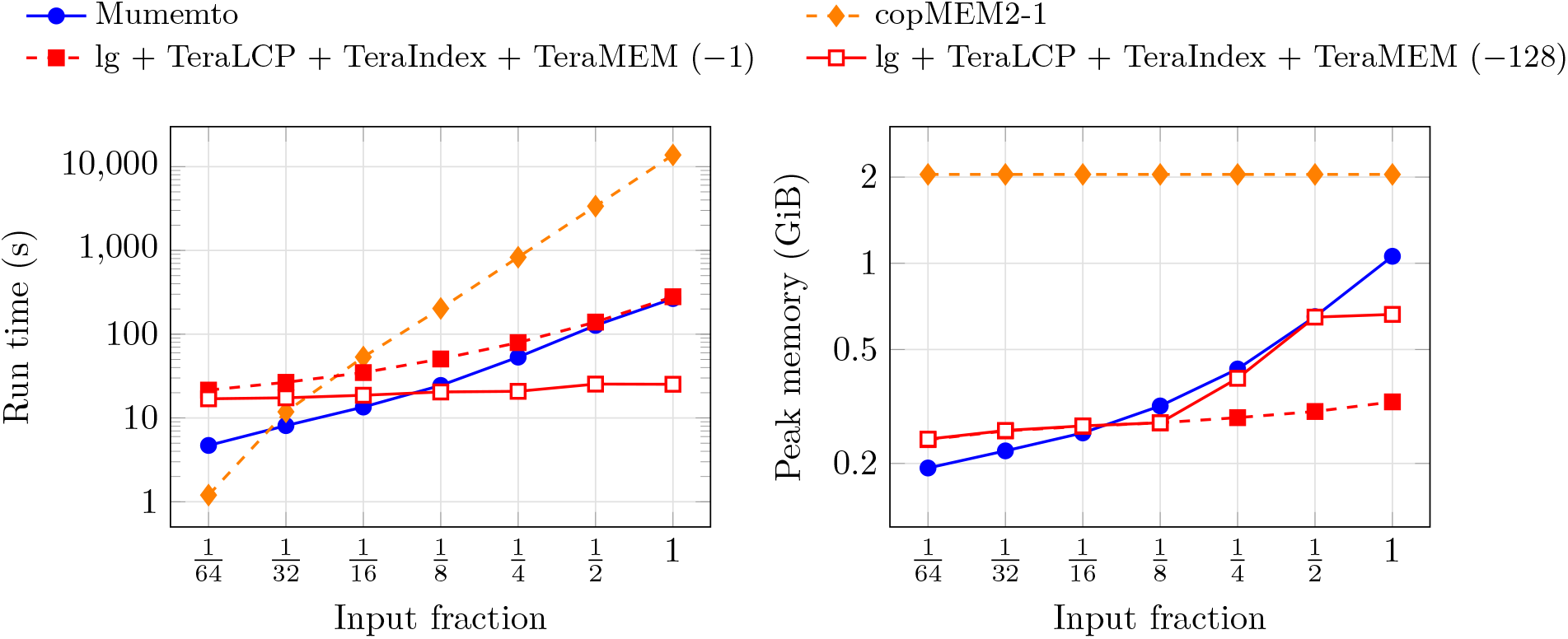
Run time and peak memory for computing MEMs from text (`mtb152`).

**Figure A4:**
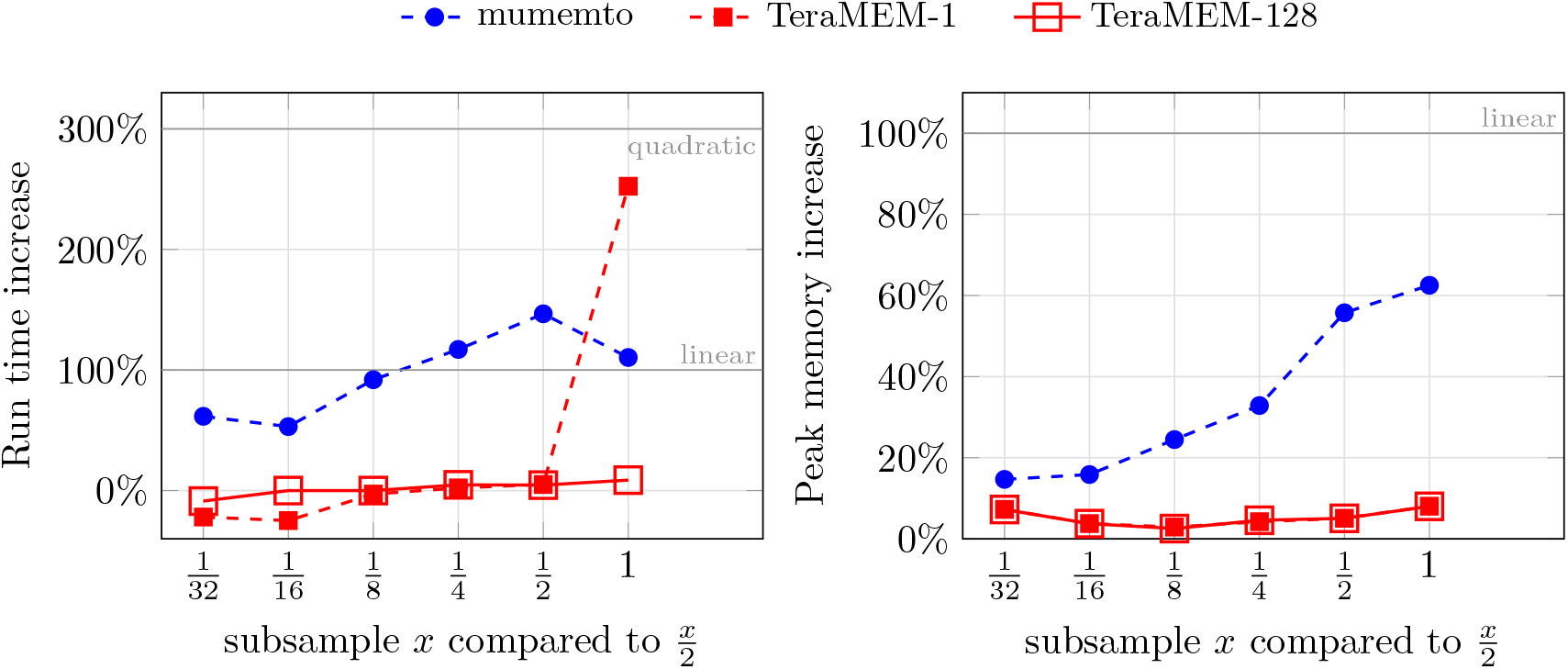
Resource requirement scaling for computing MEMs from a compressed index as datasets double in size for subsets of mtb152. Time on the left and memory on the right. X-axis is size of the twice as large dataset, y-axis is the % additional resource required relative to the subsampled dataset half the size. 300% additional resources is quadratic growth, 100% is linear, and *<* 100% is sublinear.

**Figure A5:**
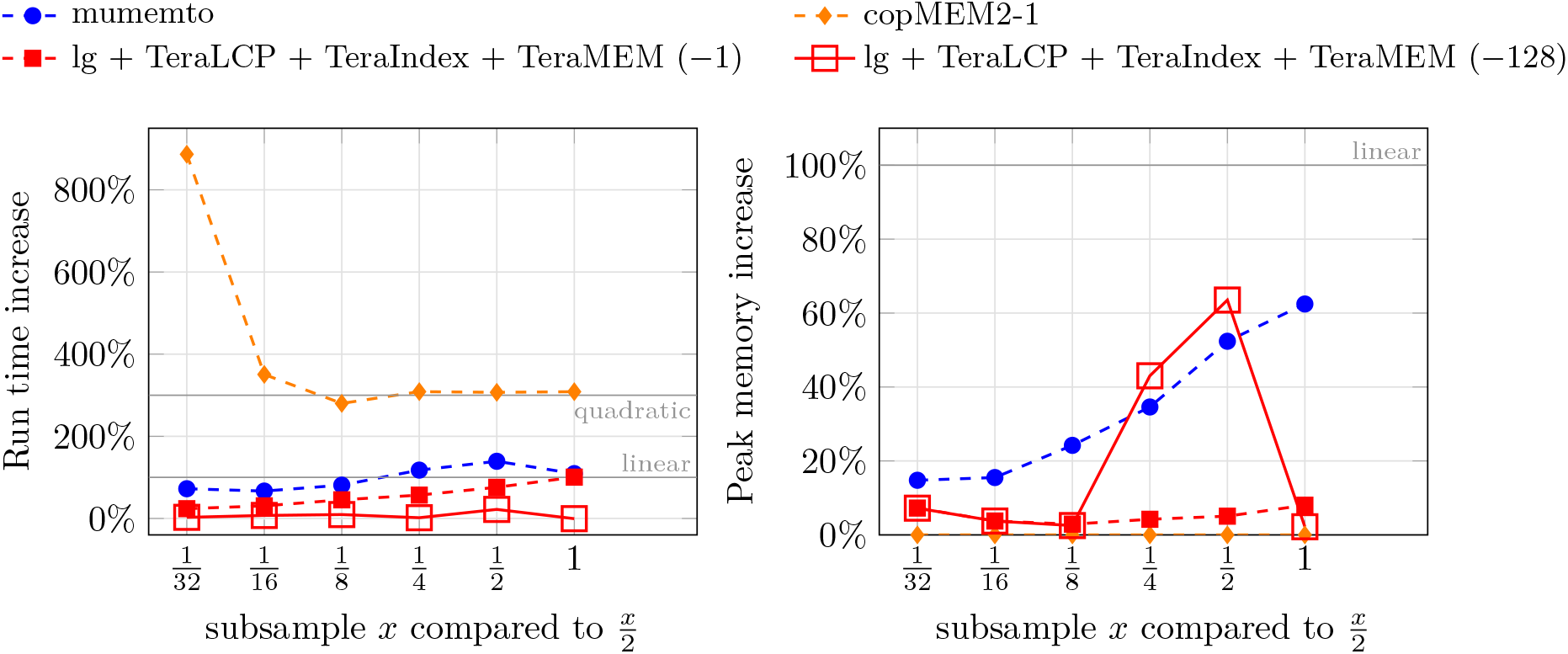
Resource requirement scaling for computing MEMs directly from the text as datasets double in size for subsets of mtb152. Time on the left and memory on the right. X-axis is size of the twice as large dataset, y-axis is the % additional resource required relative to the subsampled dataset half the size. 300% additional resources is quadratic growth, 100% is linear, and *<* 100% is sublinear.

#### D.2 humCent625

**Figure A6:**
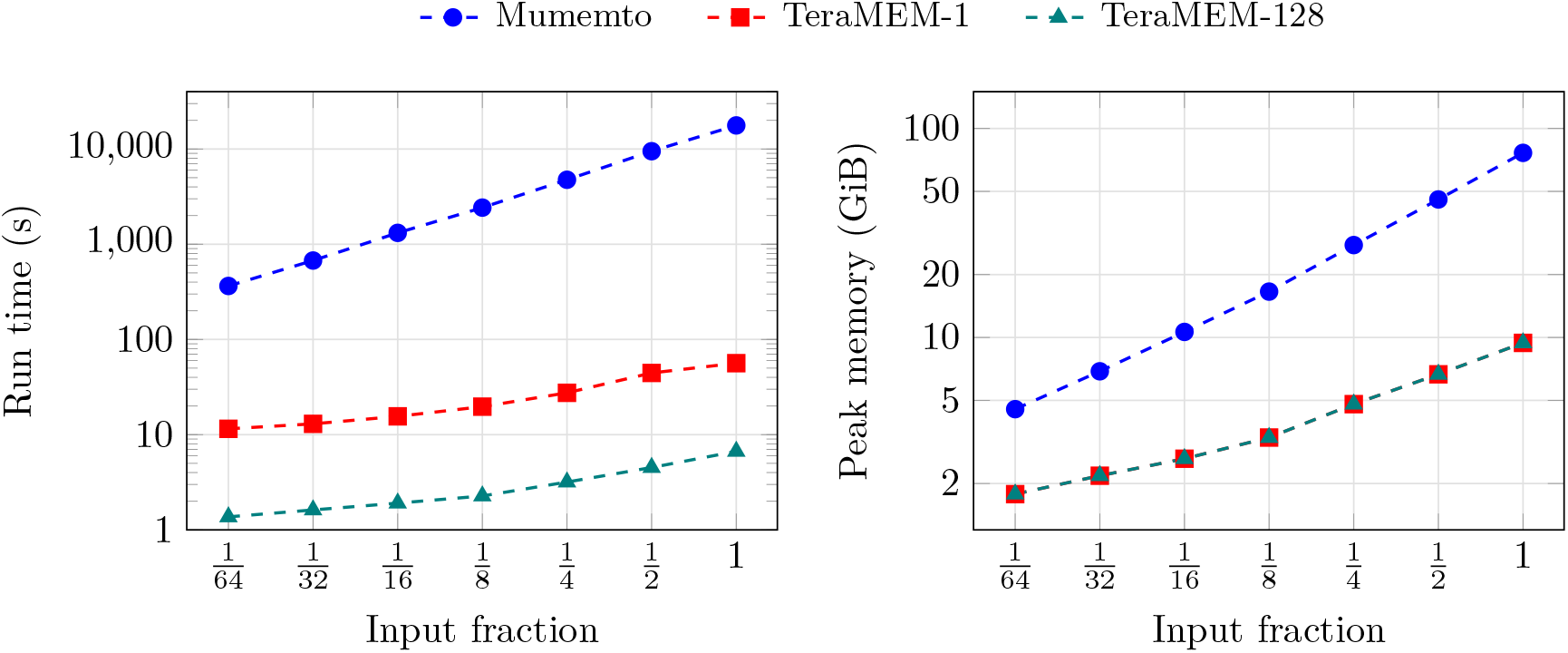
Run time and peak memory for computing MEMs from a compressed index on subsets of `humCent625`.

**Figure A7:**
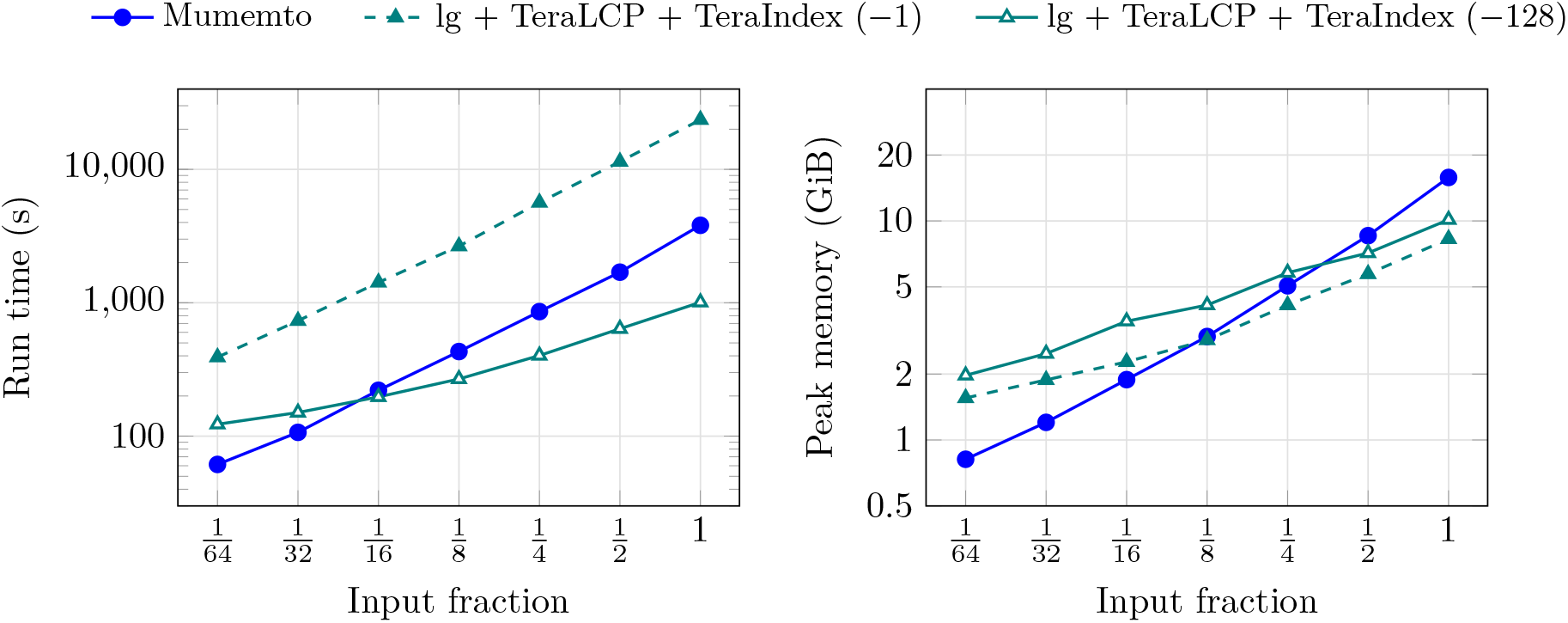
Run time and peak memory for building the compressed index input from text for subsets of `humCent625`.

**Figure A8:**
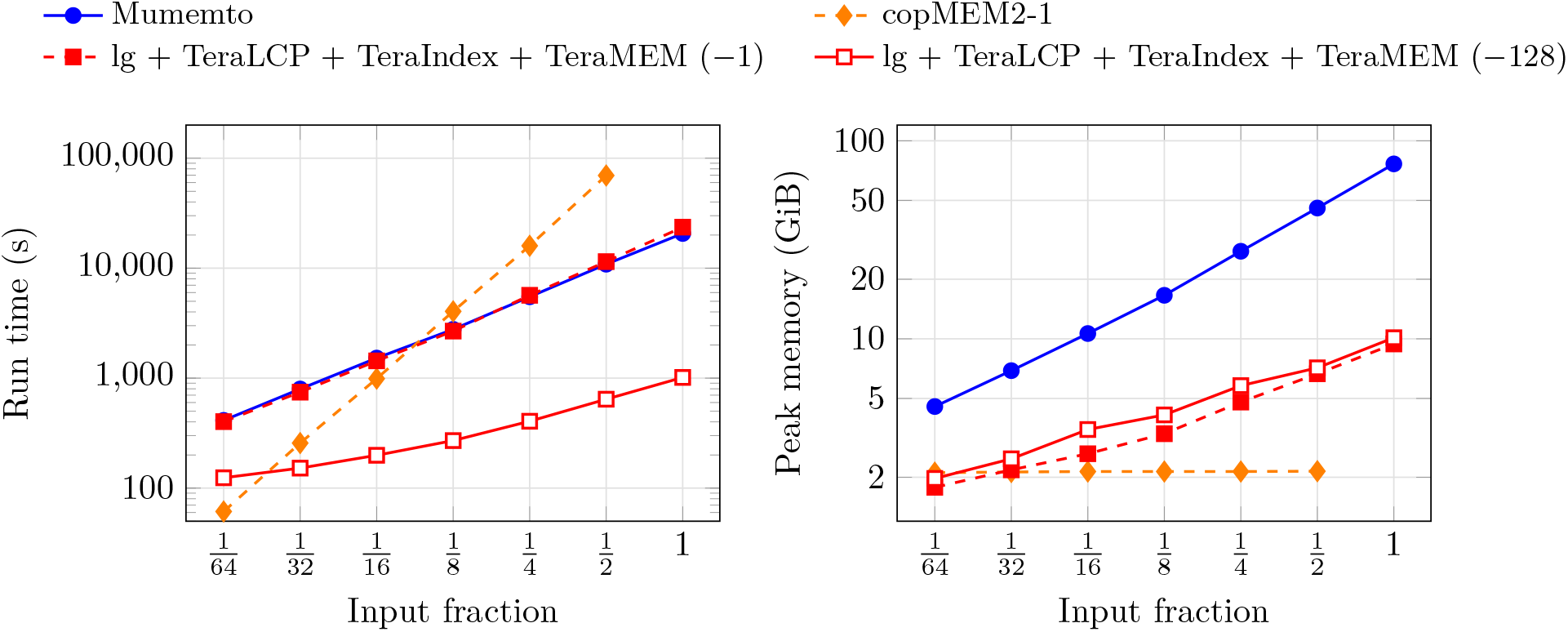
Run time and peak memory for computing MEMs from the text for subsets of `humCent625`.

**Figure A9:**
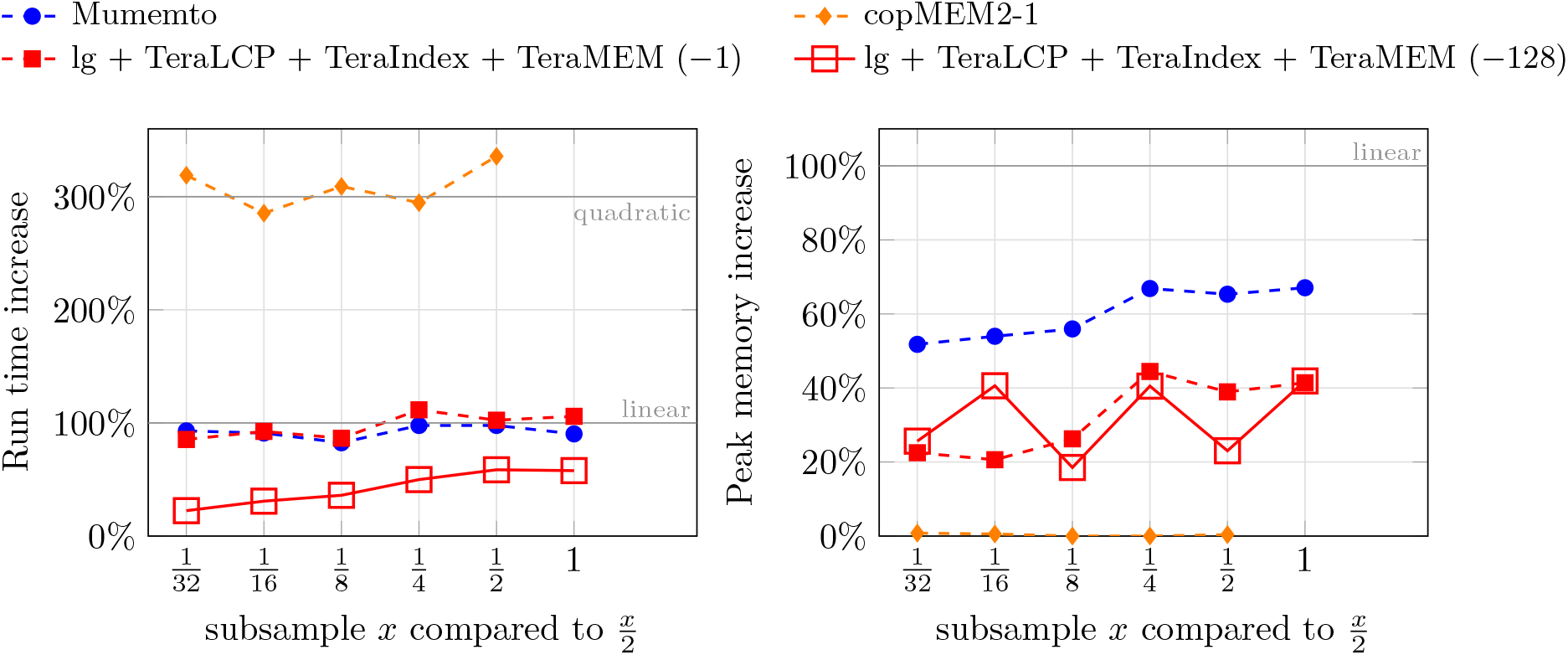
Resource requirement scaling for computing MEMs directly from the text as datasets double in size for subsets of humCent625. Time on the left and memory on the right. X-axis is size of the twice as large dataset, y-axis is the % additional resource required relative to the subsampled dataset half the size. 300% additional resources is quadratic growth, 100% is linear, and *<* 100% is sublinear.

### E Centromere Figures

#### E.1 Interchromosomal MEMs

**Figure A10:**
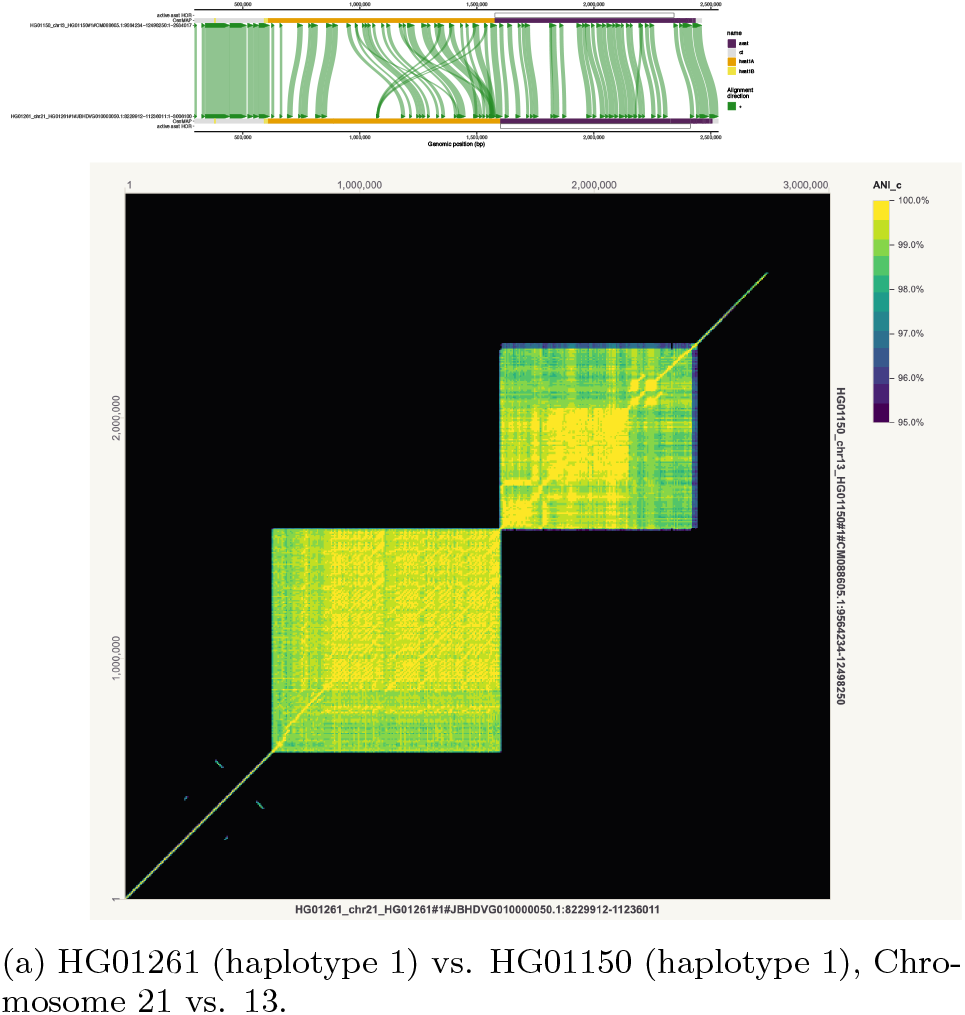
Chromosomes 13 and 21 Homology (known) [14].

#### E.2 Inversions

**Figure A11:**
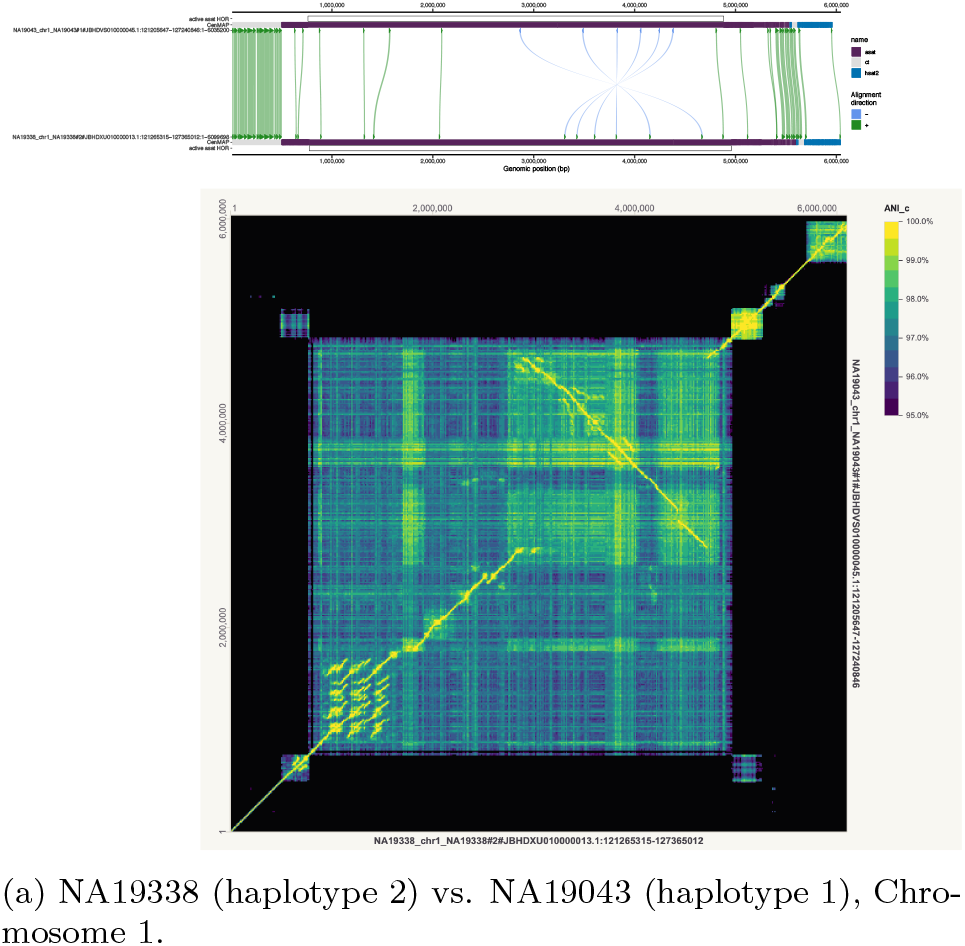
Known large inversion in Chromosome 1 [1].

**Figure A12:**
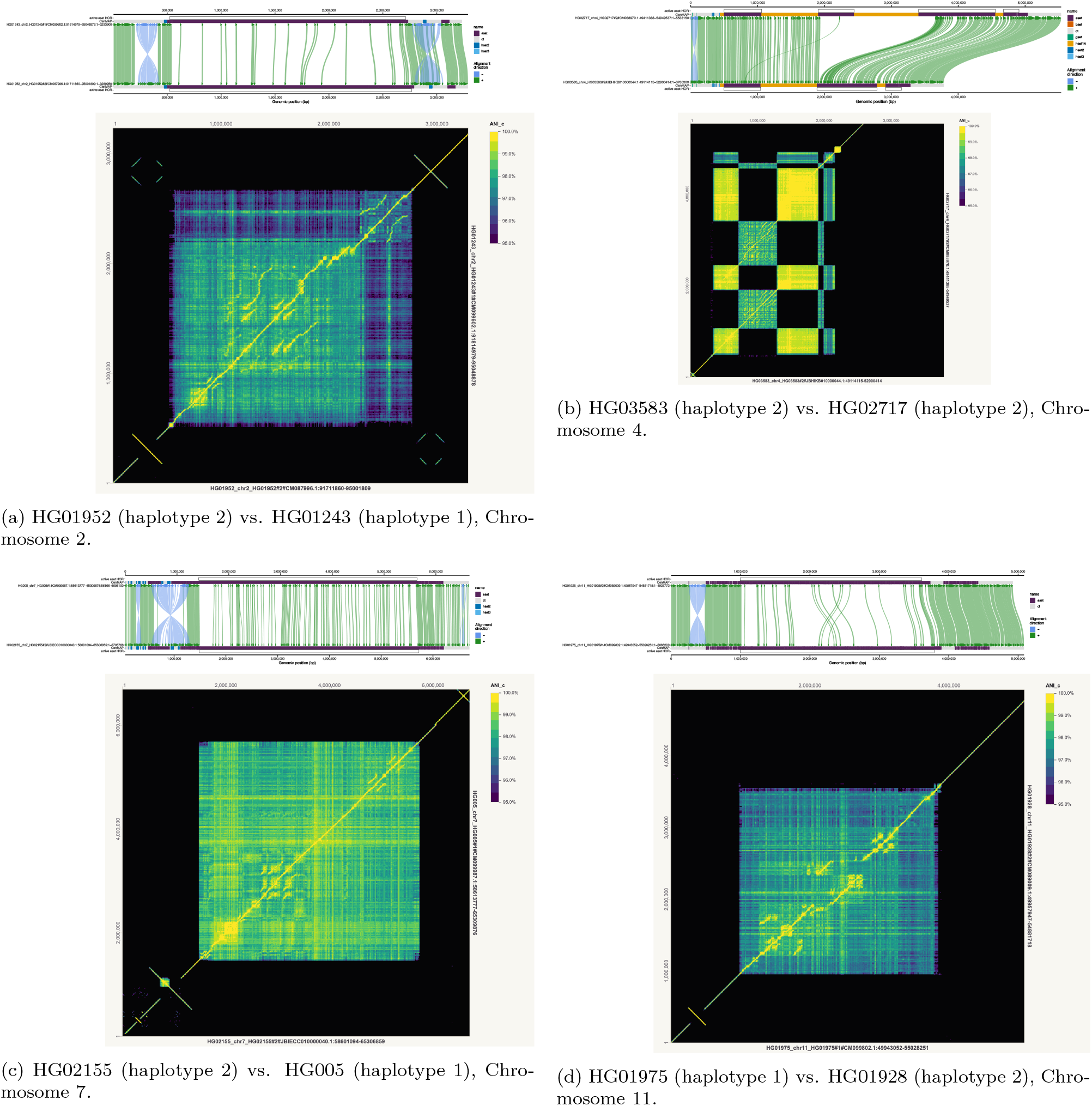
Inversions around active *α*-satellite HOR array. Large segmental duplication visible in Chromosome 4.

**Figure A13:**
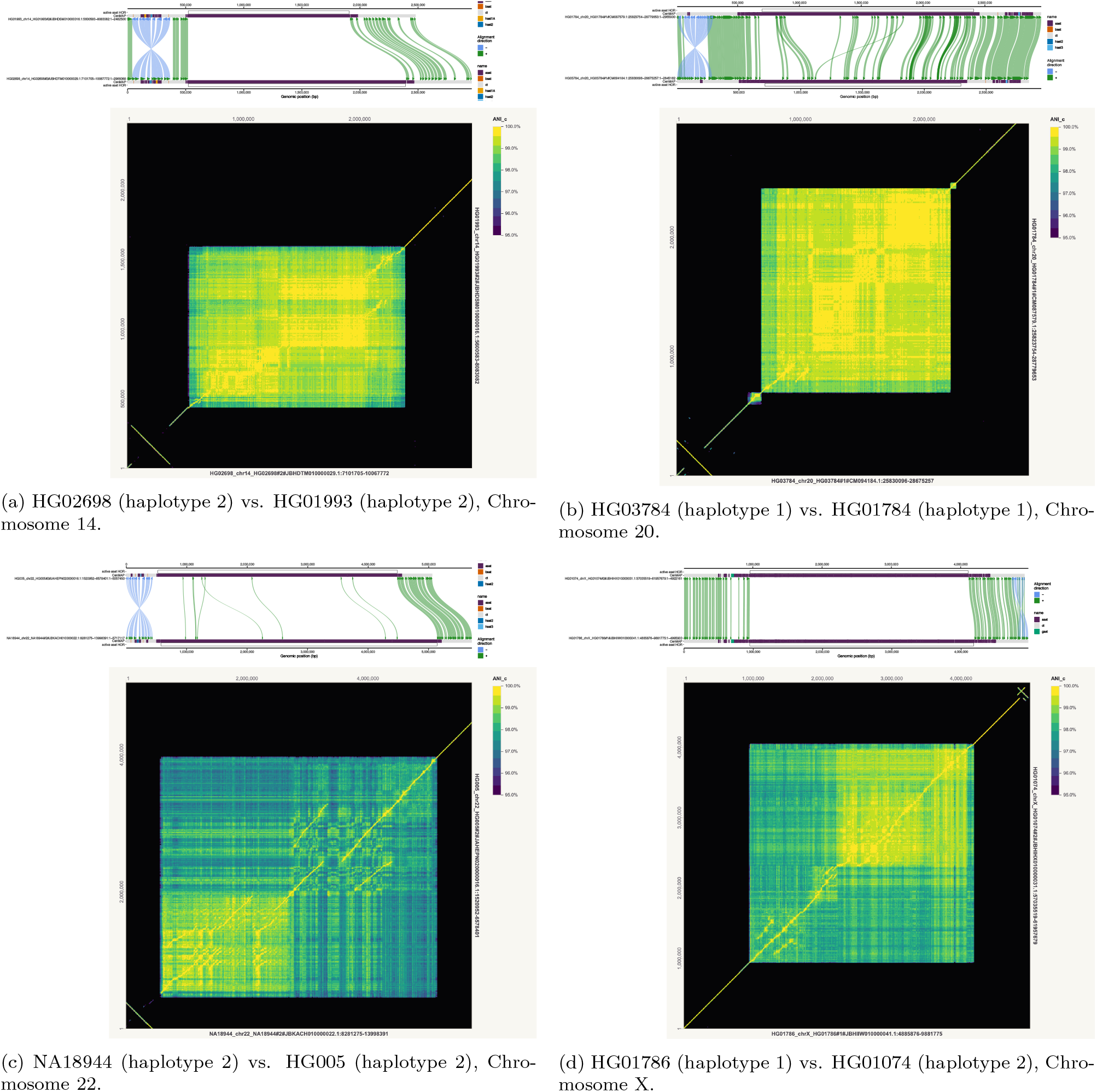
Inversions around active *α*-satellite HOR array.

#### E.3 Segmental Duplications

**Figure A14:**
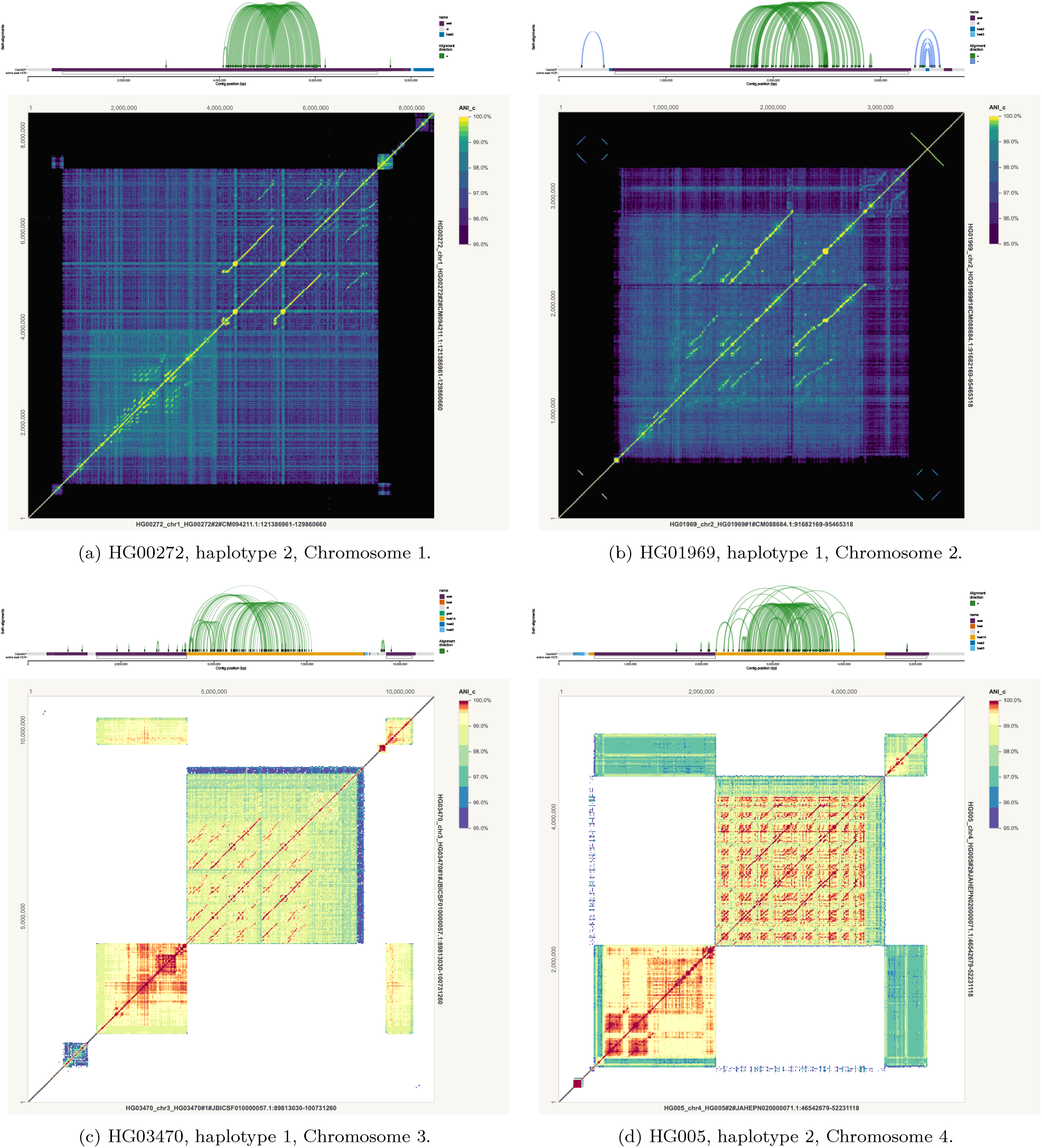
Large segmental duplications inside and outside of active *α*-satellite HOR arrays.

**Figure A15:**
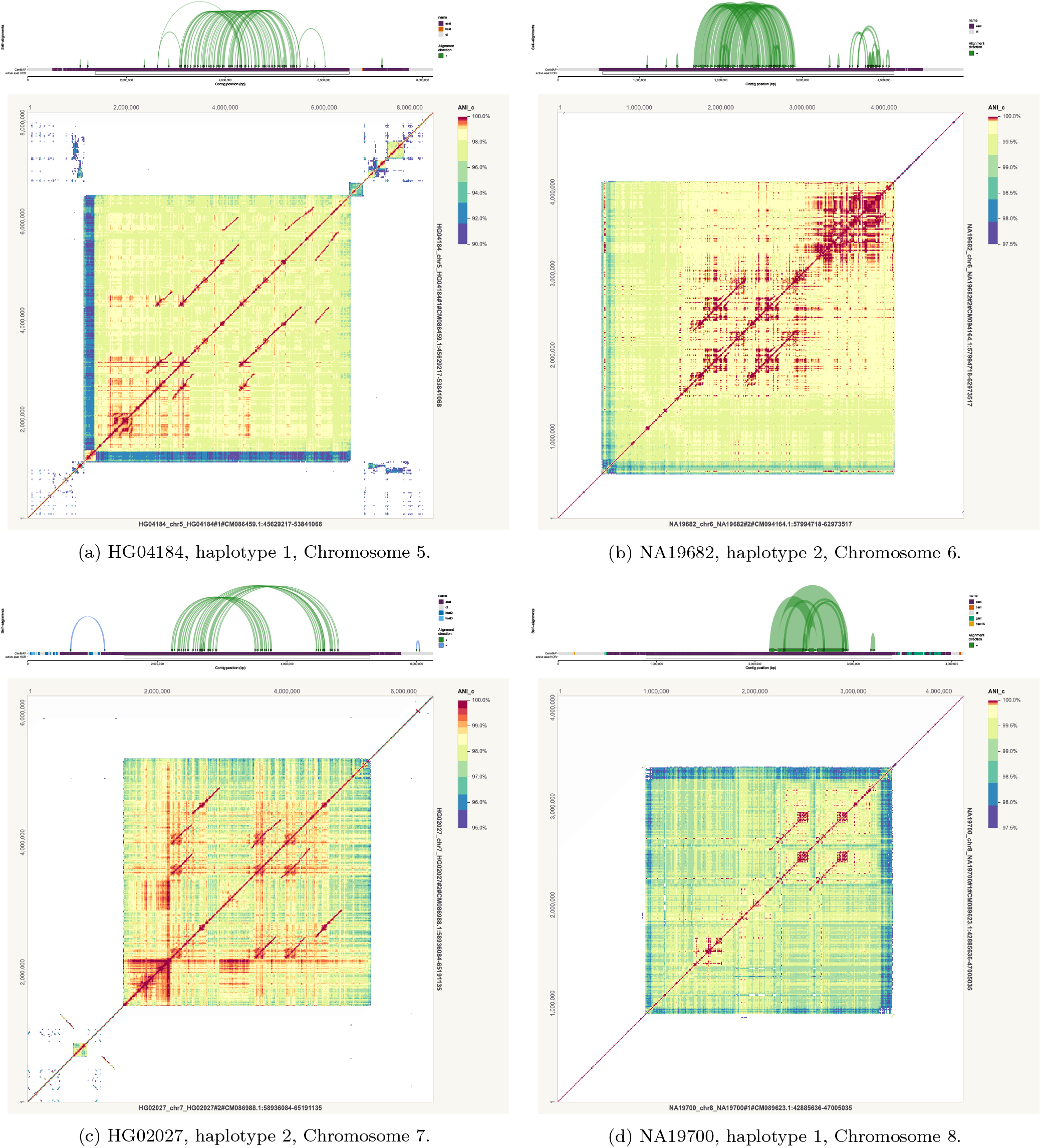
Large segmental duplications inside and outside of active *α*-satellite HOR arrays.

**Figure A16:**
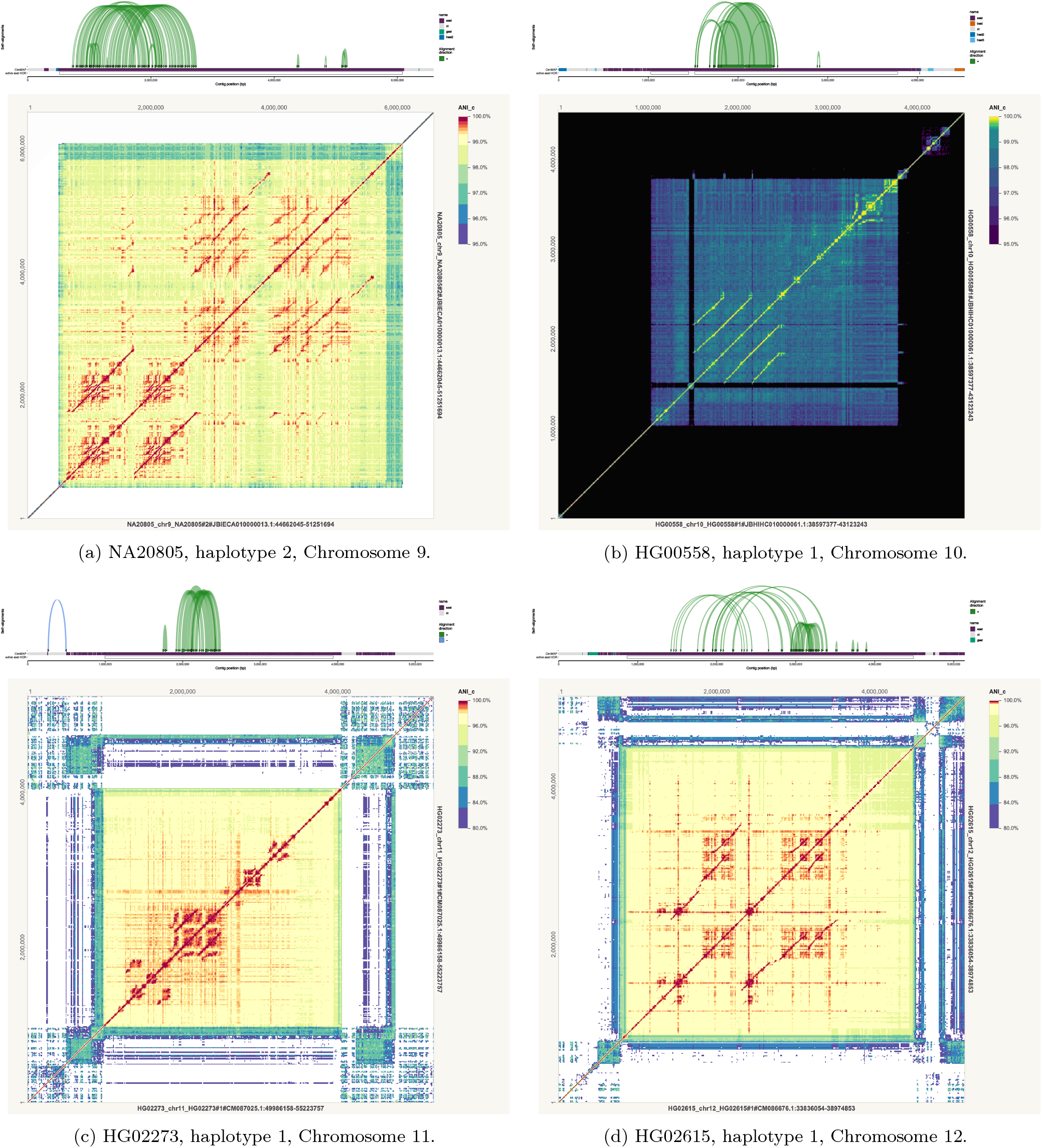
Segmental duplications in active *α*-satellite HOR arrays.

**Figure A17:**
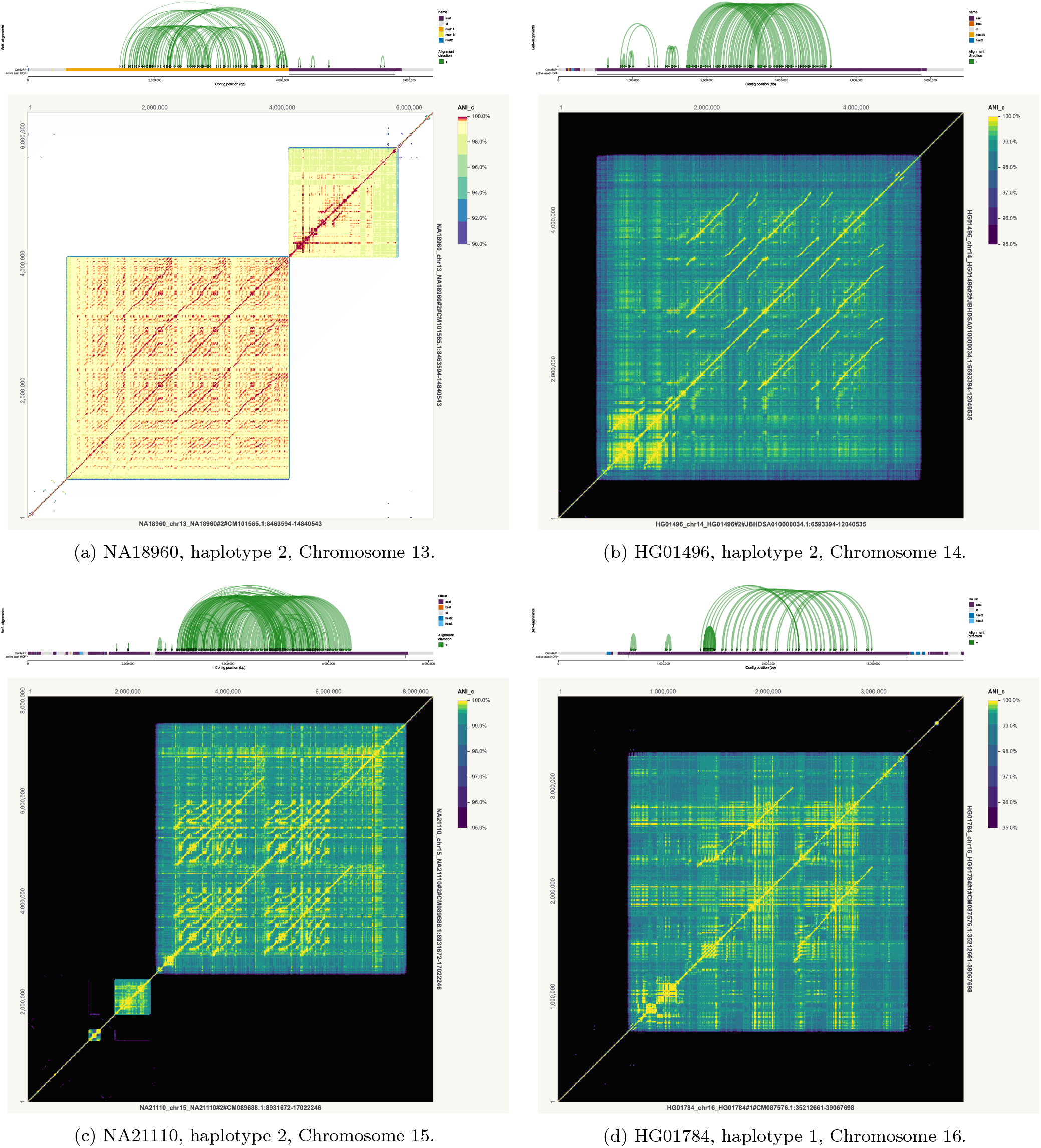
Segmental duplications inside and outside of active *α*-satellite HOR arrays.

**Figure A18:**
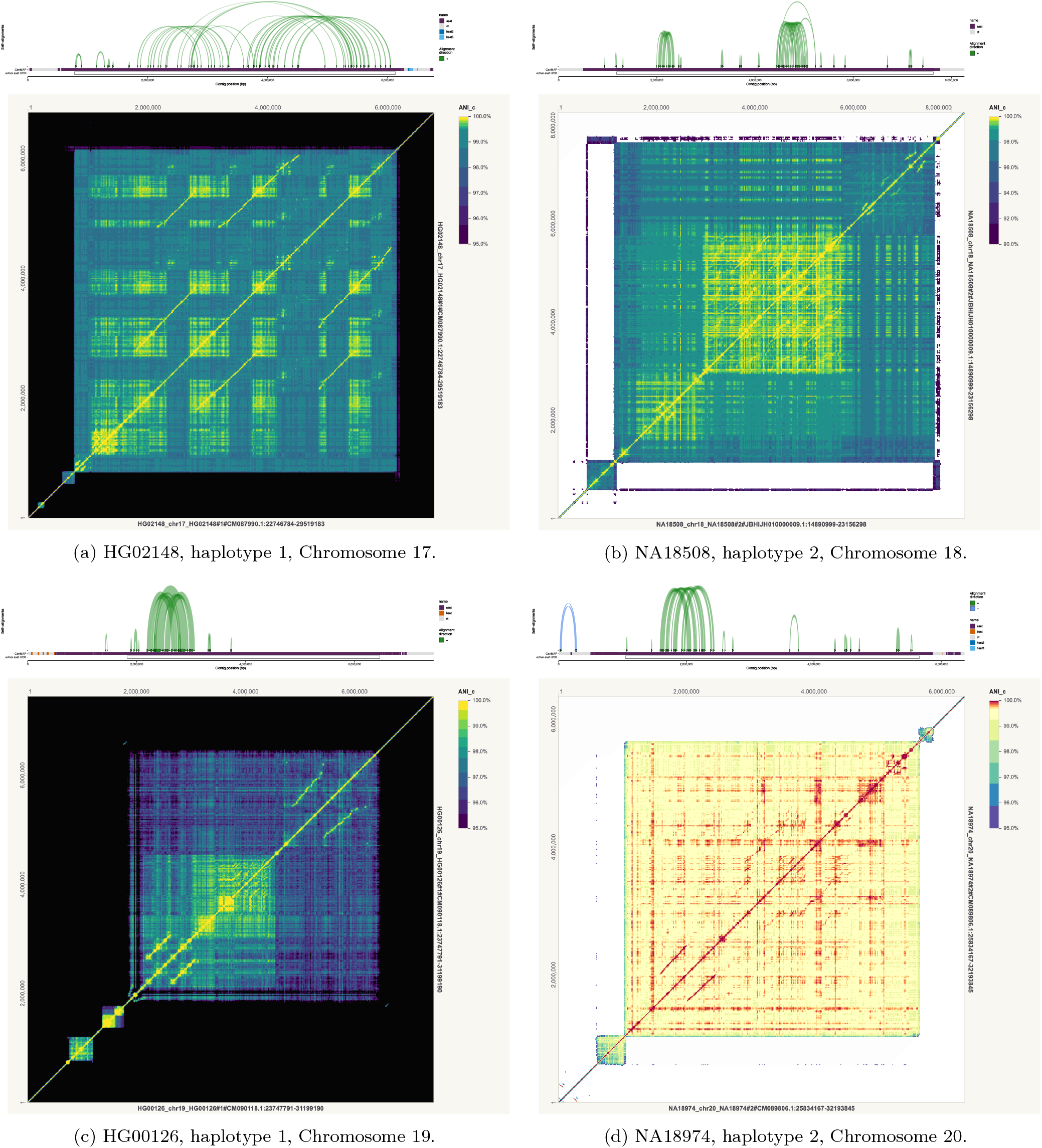
Segmental duplications in active α-satellite HOR arrays.

**Figure A19:**
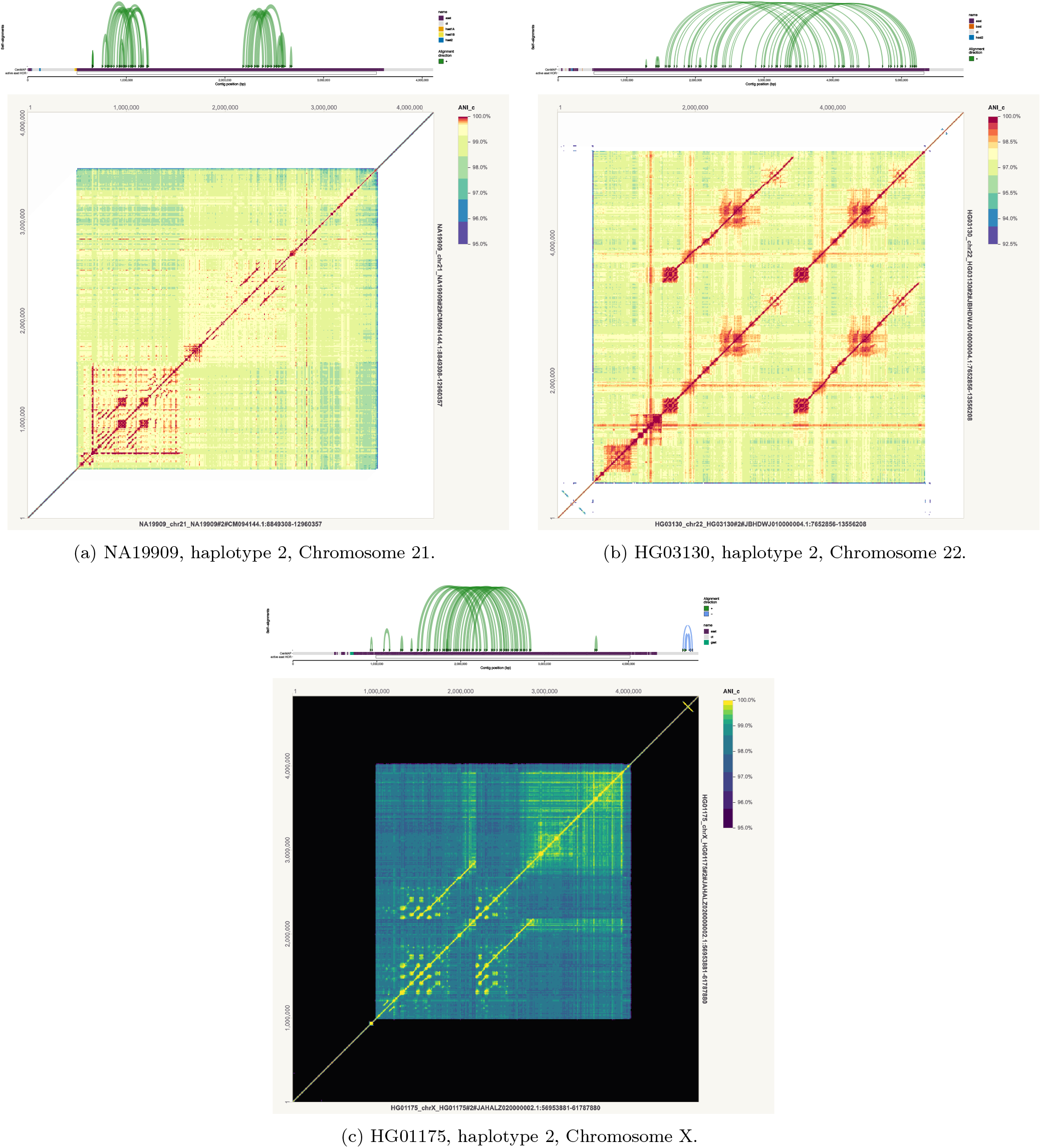
Segmental duplications in active α-satellite HOR arrays.

**Figure A20:**
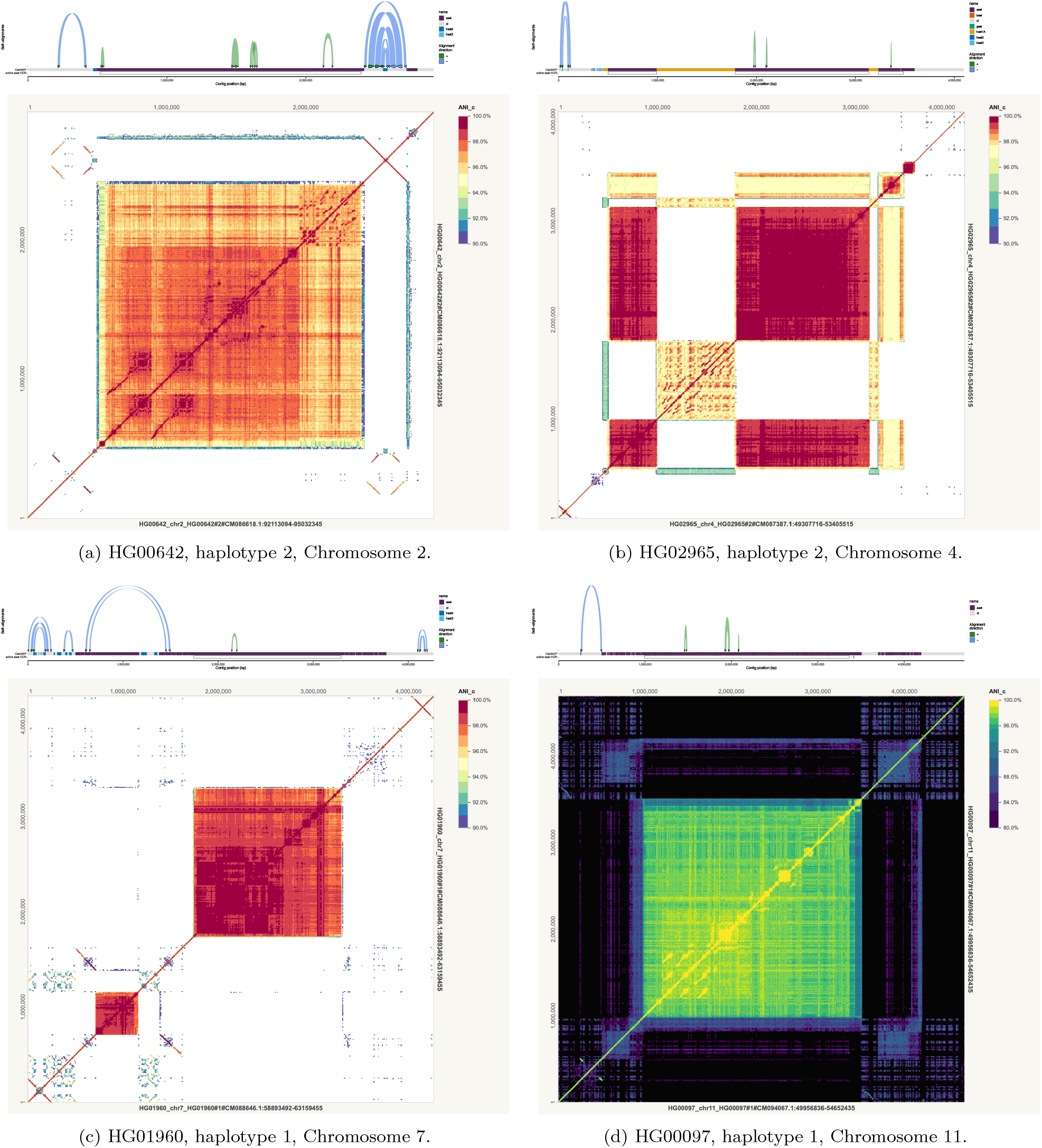
Pseudopalindromes and inverted repeats. The pseudopalindrome and inverted repeat on opposite sides of the Chromosome 2 centromere share similar sequence. Chromosome 7 has a similar dynamic.

**Figure A21:**
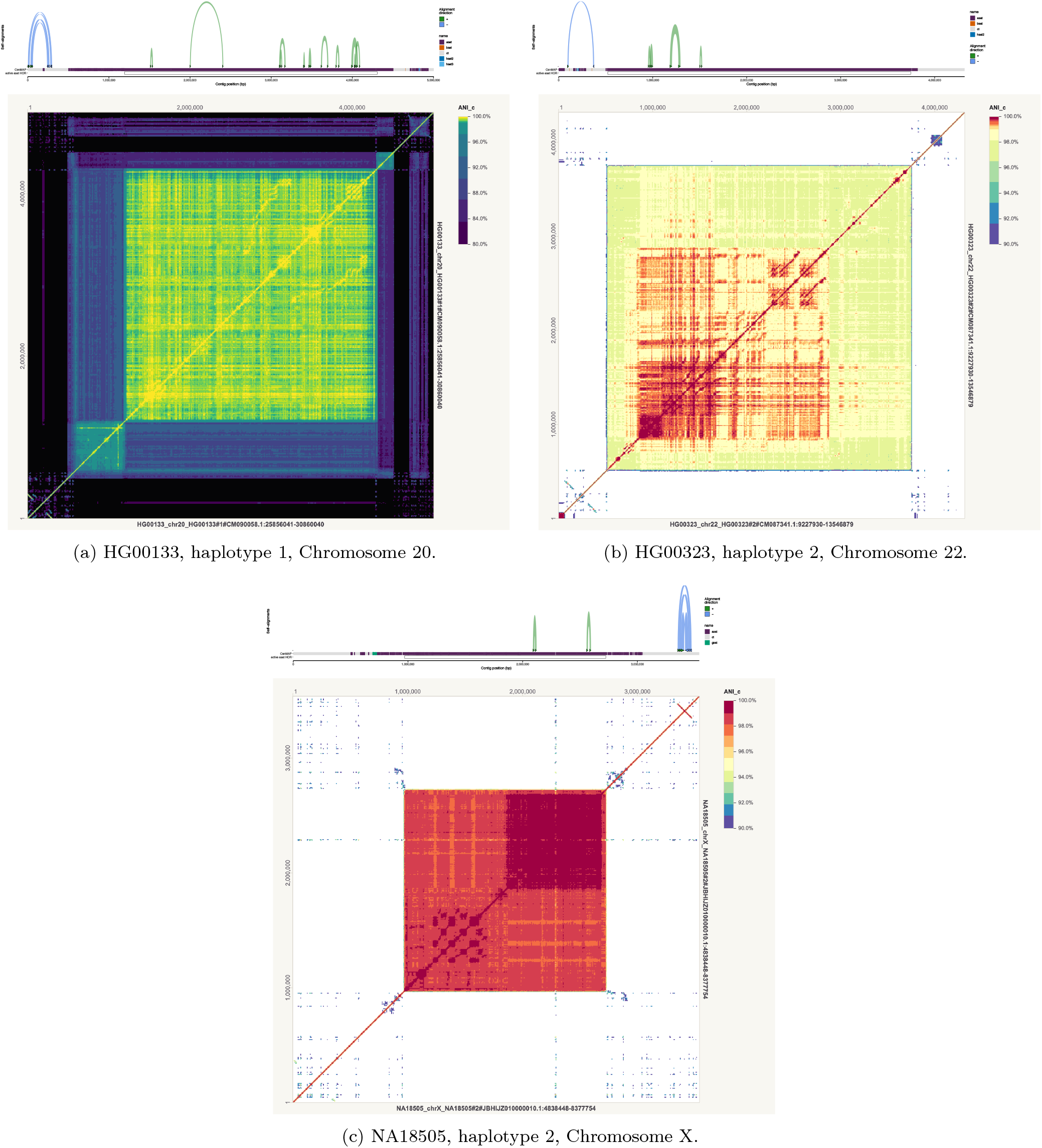
Smaller pseudopalindromes and inverted repeats.

## Footnotes

1 The original move structure description by Nishimoto and Tabei [28] differs slightly from the one provided here. In the original, some of the input intervals may be split into sub-intervals (note that all subintervals of conserved intervals are conserved intervals). Then for some position *I ∈* [0, *n −* 1], the index of the interval in this split interval sequence is required as input and the index of the interval in the split interval sequence containing *f* (*i*) is returned as output. This may be converted to our description by adding a bit vector over the split interval sequence where an index is 1 if the starting position of that interval was also a starting position of an interval in the original interval sequence. Then the interval indexes are converted between those in the split interval sequence and those in the original interval sequence through a rank or select query in constant time and *O*(|*I*|) bits of space. This doesn’t change any of the time or space complexities discussed in this paper.

## References

[1] Nicolas Altemose et al. “Complete genomic and epigenetic maps of human centromeres”. In: Science 376.6588 (2022), eabl4178. doi: 10.1126/science.abl4178. eprint: https://www.science.org/doi/pdf/10.1126/science.abl4178. URL: https://www.science.org/doi/abs/10.1126/science.abl4178.

[2] Ruben Becker et al. Compressing Suffix Trees by Path Decompositions. 2026. arXiv: 2506.14734 [cs.DS]. URL: https://arxiv.org/abs/2506.14734.

[3] Nico Bertram, Johannes Fischer, and Lukas Nalbach. “Move-r: Optimizing the r-index”. In: 22nd International Symposium on Experimental Algorithms (SEA 2024). Ed. by Leo Liberti. Vol. 301. Leibniz International Proceedings in Informatics (LIPIcs). Dagstuhl, Germany: Schloss Dagstuhl –Leibniz-Zentrum für Informatik, 2024, 1:1–1:19. ISBN: 978-3-95977-325-6. doi: 10.4230/LIPIcs.SEA.2024.1. URL: https://drops.dagstuhl.de/entities/document/10.4230/LIPIcs.SEA.2024.1.

[4] Jihun Bhak et al. “Telomere-to-Telomere Accurate and Gapless Korean Standard Reference Genome”. In: bioRxiv (2026). doi: 10.1101/2025.11.17.688257. eprint: https://www.biorxiv.org/content/early/2026/08/31/2025.11.17.688257.full.pdf. URL: https://www.biorxiv.org/content/early/2026/08/31/2025.11.17.688257.

[5] Christina Boucher et al. “Prefix-free parsing for building big BWTs”. In: Algorithms for Molecular Biology 14.1 (May 2019), p. 13. ISSN: 1748-7188. doi: 10.1186/s13015-019-0148-5. URL: 10.1186/s13015-019-0148-5.

[6] Nathaniel K. Brown, Travis Gagie, and Massimiliano Rossi. “RLBWT Tricks”. In: 20th International Symposium on Experimental Algorithms (SEA 2022). Ed. by Christian Schulz and Bora Uçar. Vol. 233. Leibniz International Proceedings in Informatics (LIPIcs). Dagstuhl, Germany: Schloss Dagstuhl –Leibniz-Zentrum für Informatik, 2022, 16:1–16:16. ISBN: 978-3-95977-251-8. doi: 10.4230/LIPIcs.SEA.2022.16. URL: https://drops.dagstuhl.de/entities/document/10.4230/LIPIcs.SEA.2022.16.

[7] Nathaniel K. Brown et al. Optimal-Time Move Structure Construction. 2026. arXiv: 2603.22147 [cs.DS]. URL: https://arxiv.org/abs/2603.22147.

[8] Yanan Chu et al. “Approaching an error-free diploid human genome using a support-based validation framework”. In: The Innovation (2026), p. 101543. ISSN: 2666-6758. doi: 10.1016/j.xinn.2026.101543. URL: https://www.sciencedirect.com/science/article/pii/S2666675826002900.

[9] S. T. Cole et al. “Deciphering the biology of Mycobacterium tuberculosis from the complete genome sequence”. In: Nature 393.6685 (June 1998), pp. 537–544. ISSN: 1476-4687. doi: 10.1038/31159. URL: 10.1038/31159.

[10] Diego Díaz-Domínguez and Leena Salmela. “Computing All-vs-All MEMs in Grammar-Compressed Text”. In: String Processing and Information Retrieval. Ed. by Franco Maria Nardini, Nadia Pisanti, and Rossano Venturini. Cham: Springer Nature Switzerland, 2023, pp. 157–170. ISBN: 978-3-031-43980-3.

[11] Richard Durbin. “A Run-Length-Compressed Skiplist Data Structure for Dynamic GBWTs Supports Time- and Space-Efficient Pangenome Operations over Syncmers”. In: Journal of Computational Biology (2026), p. 15578666261495122.

[12] Richard Durbin. “Efficient haplotype matching and storage using the positional Burrows–Wheeler transform (PBWT)”. In: Bioinformatics 30.9 (May 2014), pp. 1266–1272. ISSN: 1367-4803. doi: 10.1093/bioinformatics/btu014. eprint: https://academic.oup.com/bioinformatics/article-pdf/30/9/1266/48921891/bioinformatics_30_9_1266.pdf. URL: 10.1093/bioinformatics/btu014.

[13] Robert Edgar. “Syncmers are more sensitive than minimizers for selecting conserved k-mers in biological sequences”. In: PeerJ 9 (2021), e10805.

[14] Jordan Eizenga et al. “Pangenome alignment reveals global diversity and evolution of human centromeric regions”. In: bioRxiv (2026). doi: 10.64898/2026.09.03.749043. eprint: https://www.biorxiv.org/content/early/2026/09/04/2026.09.03.749043.full.pdf. URL: https://www.biorxiv.org/content/early/2026/09/04/2026.09.03.749043.

[15] Travis Gagie, Gonzalo Navarro, and Nicola Prezza. “Fully Functional Suffix Trees and Optimal Text Searching in BWT-Runs Bounded Space”. In: J. ACM 67.1 (Jan. 2020). ISSN: 0004-5411. doi: 10.1145/3375890. URL: 10.1145/3375890.

[16] Shenghan Gao et al. “A global view of human centromere variation and evolution”. In: Nature (July 2026). ISSN: 1476-4687. doi: 10.1038/s41586-026-10841-9. URL: 10.1038/s41586-026-10841-9.

[17] Szymon Grabowski and Wojciech Bieniecki. “copMEM2: robust and scalable maximum exact match finding”. In: Bioinformatics 39.5 (May 2023), btad313. ISSN: 1367-4811. doi: 10.1093/bioinformatics/btad313. eprint: https://academic.oup.com/bioinformatics/article-pdf/39/5/btad313/50449670/btad313.pdf. URL: 10.1093/bioinformatics/btad313.

[18] Md Helal Hossen and Daniel Gibney. “On the Size of Higher-Order Prefix-Free Parse Representations”. In: 2026 Data Compression Conference (DCC). 2026, pp. 1–9. doi: 10.1109/DCC66757.2026.00046.

[19] Emily K. Jackson et al. “Large palindromes on the primate X Chromosome are preserved by natural selection”. In: Genome Research 31.8 (2021), pp. 1337–1352. doi: 10.1101/gr.275188.120. eprint: http://genome.cshlp.org/content/genome/31/8/1337.full.pdf. URL: http://genome.cshlp.

[20] Maxat Kulmanov et al. “A reference quality, fully annotated diploid genome from a Saudi individual”. In: Scientific Data 11.1 (Nov. 2024), p. 1278. ISSN: 2052-4463. doi: 10.1038/s41597-024-04121-2. URL: 10.1038/s41597-024-04121-2.

[21] Stefan Kurtz et al. “Versatile and open software for comparing large genomes”. In: Genome Biology 5.2 (Jan. 2004), R12. ISSN: 1474-760X. doi: 10.1186/gb-2004-5-2-r12. URL: 10.1186/gb-2004-5-2-r12.

[22] Heng Li. Aligning sequence reads, clone sequences and assembly contigs with BWA-MEM. 2013. arXiv1303.3997 [q-bio.GN]. URL: https://arxiv.org/abs/1303.3997.

[23] Glennis A. Logsdon et al. “Complex genetic variation in nearly complete human genomes”. In: Nature 644.8076 (Aug. 2025), pp. 430–441. ISSN: 1476-4687. doi: 10.1038/s41586-025-09140-6. URL: 10.1038/s41586-025-09140-6.

[24] Julian K. Lucas et al. “HPRC2: A human pangenome reference with near-complete coverage of common genetic variation”. In: bioRxiv (2026). doi: 10.64898/2026.07.21.739710. eprint: https://www.biorxiv.org/content/early/2026/07/22/2026.07.21.739710.full.pdf. URL: https://www.biorxiv.org/content/early/2026/07/22/2026.07.21.739710.

[25] Maximillian G. Marin et al. “Gene conversion is a key driver of diversity hotspots in Mycobacterium tuberculosis antigens and virulence-associated loci”. In: Proceedings of the National Academy of Sciences 123.43 (2026), e2607261123. doi: 10.1073/pnas.2607261123. eprint: https://www.pnas.org/doi/pdf/10.1073/pnas.2607261123. URL: https://www.pnas.org/doi/abs/10.1073/pnas.2607261123.

[26] Nasna Nassir et al. “A draft UAE-based Arab pangenome reference”. In: Nature Communications 16.1 (July 2025), p. 6747. issn: 2041-1723. doi: 10.1038/s41467-025-61645-w. URL: 10.1038/s41467-025-61645-w.

[27] Takaaki Nishimoto, Shunsuke Kanda, and Yasuo Tabei. An Optimal-Time RLBWT Construction in BWT-runs Bounded Space. 2022. arXiv: 2202.07885 [cs.DS]. URL: https://arxiv.org/abs/2202.07885.

[28] Takaaki Nishimoto and Yasuo Tabei. “Optimal-Time Queries on BWT-Runs Compressed Indexes”. In: 48th International Colloquium on Automata, Languages, and Programming (ICALP 2021). Ed. by Nikhil Bansal, Emanuela Merelli, and James Worrell. Vol. 198. Leibniz International Proceedings in Informatics (LIPIcs). Dagstuhl, Germany: Schloss Dagstuhl –Leibniz-Zentrum für Informatik, 2021, 101:1–101:15. ISBN: 978-3-95977-195-5. doi: 10.4230/LIPIcs.ICALP.2021.101. URL: https://drops.dagstuhl.de/entities/document/10.4230/LIPIcs.ICALP.2021.101.

[29] Jannik Olbrich. “Fast and Memory-Efficient BWT Construction of Repetitive Texts Using Lyndon Grammars”. In: 33rd Annual European Symposium on Algorithms (ESA 2025). Ed. by Anne Benoit et al. Vol. 351. Leibniz International Proceedings in Informatics (LIPIcs). Dagstuhl, Germany: Schloss Dagstuhl –Leibniz-Zentrum für Informatik, 2025, 60:1–60:19. ISBN: 978-3-95977-395-9. doi: 10.4230/LIPIcs.ESA.2025.60. URL: https://drops.dagstuhl.de/entities/document/10.4230/LIPIcs.ESA.2025.60.

[30] David Porubsky et al. “Human de novo mutation rates from a four-generation pedigree reference”. In: Nature 643.8071 (July 2025), pp. 427–436. ISSN: 1476-4687. doi: 10.1038/s41586-025-08922-2. URL: 10.1038/s41586-025-08922-2.

[31] David Porubsky et al. “SVbyEye: a visual tool to characterize structural variation among whole-genome assemblies”. In: Bioinformatics 41.6 (June 2025), btaf332. ISSN: 1367-4811. doi: 10.1093/bioinformatics/btaf332. eprint: https://academic.oup.com/bioinformatics/article-pdf/41/6/btaf332/63446293/btaf332.pdf. URL: 10.1093/bioinformatics/btaf332.

[32] Arang Rhie et al. “The complete sequence of a human Y chromosome”. In: Nature 621.7978 (Sept. 2023), pp. 344–354. ISSN: 1476-4687. doi: 10.1038/s41586-023-06457-y. URL: 10.1038/s41586-023-06457-y.

[33] Ahsan Sanaullah, Degui Zhi, and Shaojie Zhang. “An Efficient Data Structure and Algorithm for Long-Match Query in Run-Length Compressed BWT”. In: 25th International Conference on Algorithms for Bioinformatics (WABI 2025). Ed. by Bron?a Brejová and Rob Patro. Vol. 344. Leibniz International Proceedings in Informatics (LIPIcs). Dagstuhl, Germany: Schloss Dagstuhl –Leibniz-Zentrum für Informatik, 2025, 17:1–17:25. isbn: 978-3-95977-386-7. doi: 10.4230/LIPIcs.WABI.2025.17. URL: https://drops.dagstuhl.de/entities/document/10.4230/LIPIcs.WABI.2025.17.

[34] Ahsan Sanaullah et al. “RLBWT-based LCP computation in compressed space for terabase-scale pangenome analysis”. In: Bioinformatics 42.Supplement 1 (July 2026), btag268. ISSN: 1367-4811. doi: 10.1093/bioinformatics/btag268. eprint: https://academic.oup.com/bioinformatics/article-pdf/42/Supplement_1/btag268/68709549/btag268.pdf. URL: 10.1093/bioinformatics/btag268.s

[35] Prasad Sarashetti et al. “A Complete Telomere-to-Telomere Diploid Reference Genome for South Asian Population”. In: bioRxiv (2025). doi: 10.1101/2025.07.12.664550. eprint: https://www.biorxiv.org/content/early/2025/09/18/2025.07.12.664550.full.pdf. URL: https://www.biorxiv.org/content/early/2025/09/18/2025.07.12.664550.

[36] Vikram S Shivakumar and Ben Langmead. “Partitioned multi-MUM finding for scalable pangenomics with MumemtoM”. In: Genome Research 36.2 (2026), pp. 397–404.

[37] Vikram S. Shivakumar and Ben Langmead. “Mumemto: efficient maximal matching across pangenomes”. In: Genome Biology 26.1 (June 2025), p. 169. ISSN: 1474-760X. doi: 10.1186/s13059-025-03644-0. URL: 10.1186/s13059-025-03644-0.

[38] Jouni Sirén et al. “Haplotype-aware graph indexes”. In: Bioinformatics 36.2 (Jan. 2020), pp. 400–407. ISSN: 1367-4803. doi: 10.1093/bioinformatics/btz575. eprint: https://academic.oup.com/bioinformatics/article-pdf/36/2/400/48990879/btz575.pdf. URL: 10.1093/bioinformatics/btz575.

[39] Alexander P Sweeten, Michael C Schatz, and Adam M Phillippy. “ModDotPlot—rapid and interactive visualization of tandem repeats”. In: Bioinformatics 40.8 (Aug. 2024), btae493. ISSN: 1367-4811. doi: 10.1093/bioinformatics/btae493. eprint: https://academic.oup.com/bioinformatics/article-pdf/40/8/btae493/58809824/btae493.pdf. URL: 10.1093/bioinformatics/btae493.

[40] Chentao Yang et al. “The complete and fully-phased diploid genome of a male Han Chinese”. In: Cell Research 33.10 (Oct. 2023), pp. 745–761. ISSN: 1748-7838. doi: 10.1038/s41422-023-00849-5. URL: 10.1038/s41422-023-00849-5.

